# Cyclic nucleotide-induced bidirectional long-term synaptic plasticity in *Drosophila* mushroom body

**DOI:** 10.1101/2023.09.28.560058

**Authors:** Daichi Yamada, Andrew M. Davidson, Toshihide Hige

## Abstract

Activation of the cAMP pathway is one of the common mechanisms underlying long-term potentiation (LTP). In the *Drosophila* mushroom body, simultaneous activation of odor-coding Kenyon cells (KCs) and reinforcement-coding dopaminergic neurons activates adenylyl cyclase in KC presynaptic terminals, which is believed to trigger synaptic plasticity underlying olfactory associative learning. However, learning induces long-term depression (LTD) at these synapses, contradicting the universal role of cAMP as a facilitator of transmission. Here, we develop a system to electrophysiologically monitor both short-term and long-term synaptic plasticity at KC output synapses and demonstrate that they are indeed an exception where activation of the cAMP/protein kinase A pathway induces LTD. Contrary to the prevailing model, our cAMP imaging finds no evidence for synergistic action of dopamine and KC activity on cAMP synthesis. Furthermore, we find that forskolin-induced cAMP increase alone is insufficient for plasticity induction; it additionally requires simultaneous KC activation to replicate the presynaptic LTD induced by pairing with dopamine. On the other hand, activation of the cGMP pathway paired with KC activation induces slowly developing LTP, proving antagonistic actions of the two second-messenger pathways predicted by behavioral study. Finally, KC subtype-specific interrogation of synapses reveals that different KC subtypes exhibit distinct plasticity duration even among synapses on the same postsynaptic neuron. Thus, our work not only revises the role of cAMP in synaptic plasticity by uncovering the unexpected convergence point of the cAMP pathway and neuronal activity, but also establishes the methods to address physiological mechanisms of synaptic plasticity in this important model.

**Abstract Figure:** Mushroom body (MB) is the olfactory learning center of the *Drosophila* brain (left). Dopamine input activates the cAMP/Protein kinase A pathway in Kenyon cells (KCs), the principal neurons of the MB. When it coincides with KC activity, it induces presynaptic long-term depression at the synapses on the MB output neuron (Top right). A subset of dopaminergic neurons is also known to release nitric oxide, which activates the cGMP pathway. When it coincides with KC activity, it induces long-term potentiation (Bottom right). Created with BioRender.com.

## Introduction

Synaptic plasticity is a fundamental mechanism of learning. *Aplysia*, the same animal that helped prove this notion for the first time, also contributed to the identification of a cAMP-dependent pathway as a key molecular basis for synaptic plasticity (Brunelli *et al*., 1976; Castellucci *et al*., 1980). Following this discovery, the cAMP/protein kinase A (PKA) pathway was found to be one of the ubiquitously important mechanisms underlying learning-related synaptic plasticity in both vertebrates and invertebrates (Kandel *et al*., 2014). In agreement with other systems, behavioral genetics studies in *Drosophila* have linked learning defects in olfactory classical conditioning to mutations of the cAMP/PKA pathway genes, such as cAMP phosphodiesterase *dunce* (Byers *et al*., 1981) and calcium/calmodulin-activated adenylyl cyclase (AC) *rutabaga* (Livingstone *et al*., 1984).

The mushroom body (MB) is the central brain area for olfactory learning in *Drosophila*. A given odor evokes reliable spiking responses in a sparse population (∼5%) of the ∼2,000 Kenyon cells (KCs), the principal neurons of the MB (Turner *et al*., 2008; Honegger *et al*., 2011). KCs form dense axon bundles, constituting the MB lobes, where they synapse on their main postsynaptic partners, MB output neurons (MBONs). In the MB lobes, KCs also receive dense inputs from the dopaminergic neurons (DANs), which, depending on cell types, convey either reward or punishment signals during conditioning (Schwaerzel *et al*., 2003; Liu *et al*., 2012; Burke *et al*., 2012; Aso & Rubin, 2016). Thus, olfactory and reinforcement signals converge at the KC axons. This notion is consistent with the fact that memory defects of the mutants of a G_s_-linked dopamine receptor *Dop1R1* (Kim *et al*., 2007; Qin *et al*., 2012) and *rutabaga* (McGuire *et al*., 2003; Blum *et al*., 2009) can be fully rescued by expressing the corresponding functional proteins in KCs. These results led to the prevailing working model that Rutabaga AC in the KC axons acts as a coincidence detector of olfactory and reinforcement signals, represented by calcium influx and dopamine input, respectively, and the resulting increase in cAMP level induces presynaptic plasticity at KC-MBON synapses (Heisenberg, 2003). In support of this model, coactivation of KCs and dopamine receptors synergistically activates cAMP/PKA pathway in KC axons (Tomchik & Davis, 2009; Gervasi *et al*., 2010; Handler *et al*., 2019; but see also Abe *et al*., 2023).

In general, action of cAMP on synaptic transmission is excitatory. cAMP increase virtually always results in potentiation of synapses in both vertebrates and invertebrates. The examples range from synaptic facilitation of the siphon sensory neurons in *Aplysia* (Goldsmith & Abrams, 1991) to long-term potentiation (LTP) in rodent hippocampus (Huang *et al*., 1994) and cerebellum (Salin *et al*., 1996). Conversely, decrease in cAMP underlies multiple forms of long-term depression (LTD) (Tzounopoulos *et al*., 1998; Chevaleyre *et al*., 2007). This positive relationship between cAMP level and synaptic strength also seems to apply to *Drosophila* as elevated presynaptic cAMP level mediates post-tetanic synaptic potentiation at the neuromuscular junction (Kuromi & Kidokoro, 2000), which was impaired in *dunce* and *rutabaga* mutants (Zhong & Wu, 1991). Furthermore, multiple pioneering studies of either the early (Wang *et al*., 2008) or late phase (Yu *et al*., 2006; Akalal *et al*., 2010) of memory traces induced by olfactory learning reported potentiation of odor-evoked calcium activity in the KC axons.

Given this historic background, it was rather unexpected that pairing of odor presentation and optogenetic activation of DANs induced robust LTD at KC-to-MBON synapses (Hige *et al*., 2015). However, LTD but not LTP fits the circuit logic of the MB. Anatomically, a given cell type of MBONs has partner cell types of DANs, and they show matching innervation patterns in the MB lobes (Aso *et al*., 2014b). In general, activation of MBONs signals the valence that is opposite to the one signaled by the partner DANs (Aso *et al*., 2014a; Owald *et al*., 2015; Aso & Rubin, 2016). Thus, punishment-encoding DANs can induce LTD in approach-directing MBONs during aversive learning. Although numerous other studies now support or confirm that odor-specific depression in MBON responses underlies olfactory learning (Séjourné *et al*., 2011; Owald *et al*., 2015; Cohn *et al*., 2015; Perisse *et al*., 2016; Berry *et al*., 2018; Felsenberg *et al*., 2018; Handler *et al*., 2019; Awata *et al*., 2019; Zhang *et al*., 2019; McCurdy *et al*., 2021; Hancock *et al*., 2022; Schnitzer *et al*., 2022; Noyes & Davis, 2023; Zeng *et al*., 2023), there has been no direct evidence that *Drosophila* MB is an exception where cAMP-induced synaptic plasticity is depression rather than potentiation. Providing such evidence is the main objective of this study. cAMP signaling is not the only second messenger system implicated in learning-related plasticity in the *Drosophila* MB. A subset of the MB-projecting DANs releases nitric oxide (NO) as a cotransmitter (Aso *et al*., 2019). Behavioral evidence suggests that NO acts on KC axons to induce plasticity that is antagonistic to dopamine-induced LTD via activation of soluble guanylyl cyclase (sGC) (Aso *et al*., 2019), although there has been no physiological evidence for it.

Unlike cAMP, the sign of synaptic plasticity induced by cGMP varies among systems and studies. While cGMP increase predominantly induces presynaptic LTP in the hippocampus (Arancio *et al*., 1995) and hyperexcitability of sensory neurons in *Aplysia* (Lewin & Walters, 1999), it also induces LTD in the hippocampus (Reyes-Harde *et al*., 1999), cerebellum (Shibuki & Okada, 1991; Lev-Ram *et al*., 1997) and corticostriatal synapses (Calabresi *et al*., 1999). At the *Drosophila* neuromuscular junction, cGMP exerts an excitatory (Wildemann & Bicker, 1999) or no effect (Caplan *et al*., 2013). Furthermore, NO-dependent modulation of the *Drosophila* neuromuscular junction also involves cGMP-independent, S-nitrosylation of proteins (Robinson *et al*., 2018). Thus, it is important to determine the role of cGMP in KC-to-MBON synaptic plasticity.

In this study, we developed an *ex vivo* system to test physiological and pharmacological properties of the synaptic plasticity at KC-to-MBON synapses. In this system, we made whole-cell recordings from a target MBON to monitor the EPSCs evoked by optogenetic stimulation of a small subset of KCs, while focally injecting various reagents to the MB lobe at the dendritic region of the MBON to induce or inhibit long-term plasticity. This system also allowed us to monitor short-term synaptic plasticity by delivering paired-pulse stimulation to test the involvement of presynaptic factors in synaptic changes (Zucker & Regehr, 2002). We show that pairing of KC activation and dopamine injection induces LTD accompanied by an increase in paired-pulse ratio (PPR). Unexpectedly, however, activation of AC by forskolin alone was insufficient to induce qualitatively similar LTD; it required simultaneous KC activation in addition to forskolin application. Pairing of pharmacological activation of sGC and KC activation induced LTP accompanied by a decrease in PPR. Our system also allowed for subtype-specific activation of KCs that synapse on the same MBON and revealed distinct durations of synaptic plasticity between different KC subtypes. Thus, our work not only revises the role of cAMP in synaptic plasticity by revealing an unexpected convergence point of the cAMP pathway and neuronal activity, but also establishes the methods to address physiological mechanisms of synaptic plasticity.

## Methods

### Flies

All fly stocks were maintained at room temperature on conventional cornmeal-based medium. However, in most cases, we kept the final crosses for experiments in the dark at 18 °C to minimize the potential phototoxicity on KCs expressing CsChrimson. Flies were selected for desired genotypes on the day of eclosion, transferred to all-trans-retinal food (0.5 mM) and used for experiments after 48-72 hours. For experiments to record γ KC-to-MBON-γ1pedc synaptic currents (Figs. 1-3, 6 and 7), we used *nSyb-IVS-phiC31 attp18/w; 20XUAS-SPARC2-S-Syn21-CsChrimson::tdTomato-3.1 CR-P40/R12G04-LexA attP40; MB623C/pJFRC57-13XLexAop2-IVS-GFP-p10 VK00005*. For α/β KC-to-MBON-γ1pedc synapses (Figs. 8-10), we used *MB008C* instead of *MB623C*. For cAMP imaging (Fig. 5), we used *nSyb-IVS-phiC31 attp18/w; 20XUAS-SPARC2-S-Syn21-CsChrimson::tdTomato-3.1 CR-P40/+; MB623C/UAS-G-Flamp1 attp2*. *nSyb-IVS-phiC31 attp18* and *20XUAS-SPARC2-S-Syn21-CsChrimson::tdTomato-3.1 CR-P40* flies were obtained from the Bloomington Drosophila Stock Center. MB623C and G-Flamp1 flies were gifted from Yoshinori Aso (Janelia, HHMI) and Wanhe Li (Texas A&M), respectively.

**Figure 1.**
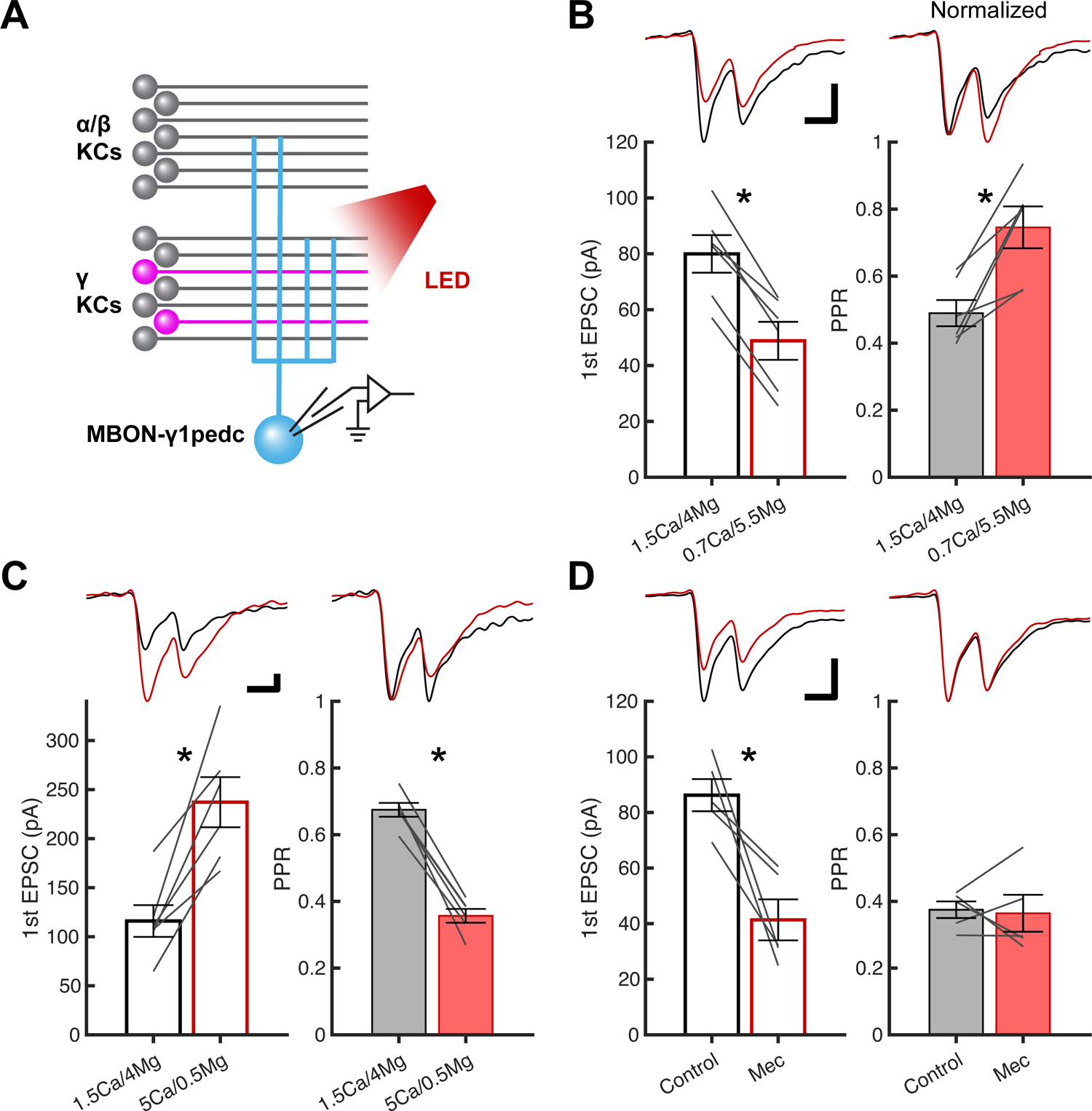
Optogenetic assessment of short-term synaptic plasticity at KC-to-MBON synapses A, a schematic of the experiments. Optogenetically evoked EPSCs were measured at γ KC-to-MBON-γ1pedc synapses by whole-cell voltage-clamp recordings from MBON-γ1pedc. Short-term plasticity induced by paired pulses (pulse width, 1 ms; interval, 400 ms) was monitored while changing the extracellular concentrations of divalent cations or partially blocking postsynaptic ionotropic receptors. B, changing the extracellular calcium/magnesium concentrations from 1.5/4 mM to 0.7/5.5 mM decreased the first EPSC amplitude (left, mean ± SEM; n = 6, p < 10^−4^, paired t-test) while increasing the PPR (right, p = 0.00658). Gray lines indicate data from individual cells. Upper left traces show overlaid representative EPSCs before (black) and after (red) changing the extracellular saline. Horizontal and vertical scale bars in this and the other panels indicate 300 ms and 30 pA, respectively. Upper right traces show the same EPSCs normalized with the first EPSC amplitude. Asterisks denote p < 0.05. C, changing the extracellular calcium/magnesium concentrations to 5/0.5 mM increased the first EPSC amplitude (left; n = 6, p = 0.00778, paired t-test) while decreasing the PPR (right; p < 10^−4^). D, bath application of mecamylamine (Mec; 10 µM), a non-competitive antagonist of the nicotinic acetylcholine receptors, reduced the first EPSC amplitude (left; n = 5, p = 0.0144, paired t-test) without affecting the PPR (right, p = 0.854).

**Figure 2.**
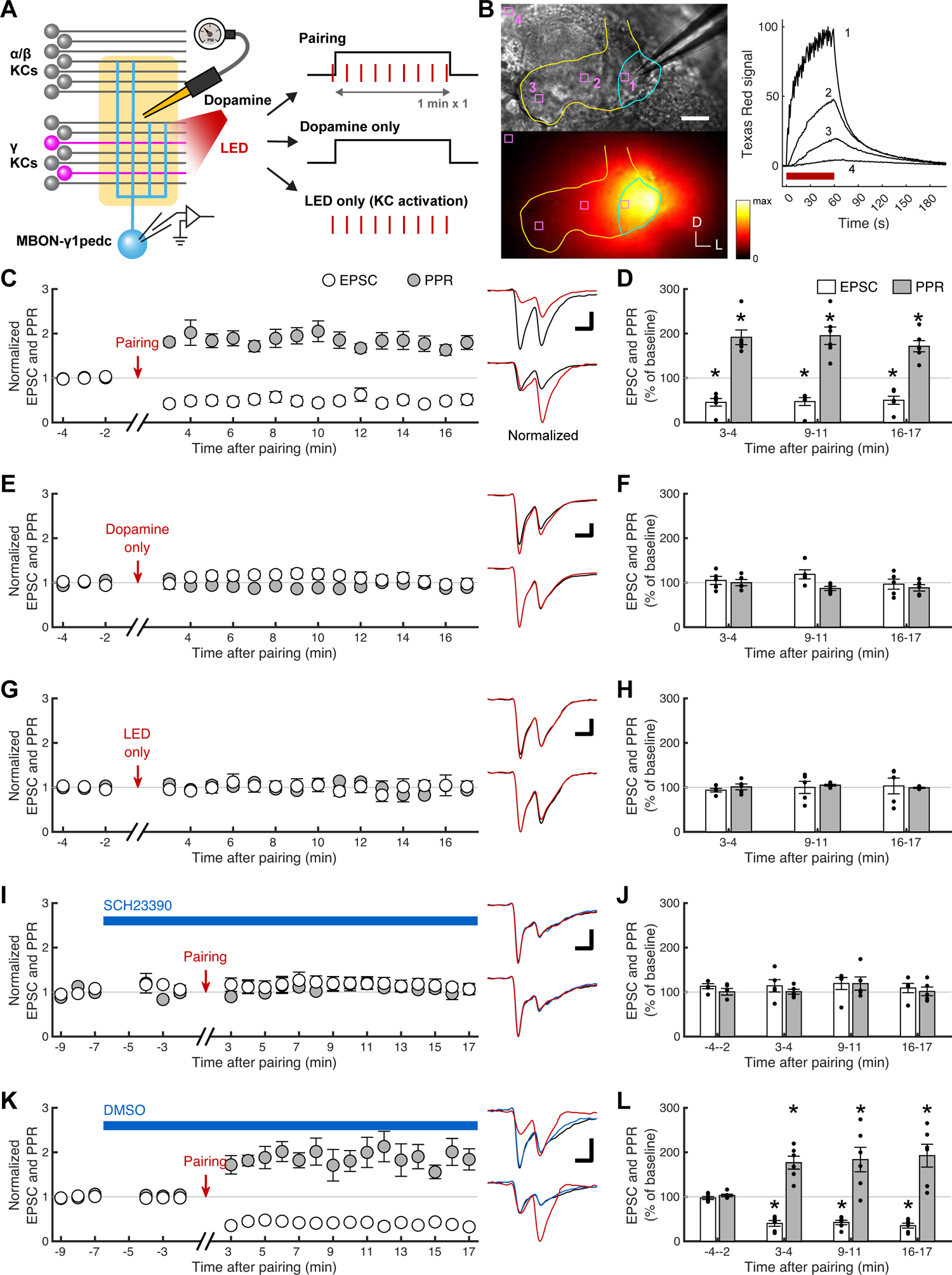
Pairing **γ** KC activation with focal dopamine application induces presynaptic LTD via D_1_-like dopamine receptors A, a schematic of the experiments. Dopamine (1 mM) was focally applied to the dendritic region of the MBON-γ1pedc via an injection pipette while measuring optogenetically evoked γ KC-to-MBON-γ1pedc EPSCs. See Methods for detailed parameters for injections and recordings. B, a representative image (lower left) to show the spread of the fluorescent signal of Texas Red dextran, which was infused with dopamine in the injection pipette, after 1-min injection. Upper left image shows a widefield view of the same sample. The γ lobe and part of the vertical lobes are outlined by yellow. Light blue line indicates the approximate location of the γ1pedc region. Signals measured in four regions of interest (light magenta squares) were plotted on the right. Horizontal red bar denotes the timing of the injection. D: dorsal, L: lateral, scale bar: 20 µm. C, first EPSC amplitudes (open circles, mean ± SEM; n = 6) and PPRs (filled circles) plotted against time after the end of 1-min pairing of γ KC activation and dopamine injection. The data were normalized to the average of a 3-min baseline recorded before pairing. Upper right traces show overlaid representative EPSCs sampled before (at −2 min; black) and after (at 3 min; red) pairing. Horizontal and vertical scale bars in this and the other panels indicate 300 ms and 30 pA, respectively. Lower right traces show the same EPSCs normalized with the first EPSC amplitude. D, quantification of the data shown in C at early, middle and late periods after pairing (mean ± SEM). Black dots indicate data from individual cells. First EPSC amplitudes (open bars) and PPRs (filled bars) showed depression and an increase, respectively, at all three time points. P values for EPSCs are (from left to right) < 10^−3^, < 10^−3^ and < 10^−3^ (Dunnett’s multiple comparisons test following repeated measures one-way ANOVA, p < 10^−3^), and for PPRs, < 10^−3^, < 10^−4^ and < 10^−3^ (repeated measures one-way ANOVA, p < 10^−4^). E, same as C, but KC activation was omitted during pairing (n = 5). F, quantification of the data shown in E. 1-min dopamine injection alone affected neither first EPSC amplitudes (p = 0.372, repeated measures one-way ANOVA) nor PPRs (p = 0.160). G, same as C, but dopamine application was omitted during pairing (n = 5). H, quantification of the data shown in G. 1-min KC activation alone affected neither first EPSC amplitudes (p = 0.632, repeated measures one-way ANOVA) nor PPRs (p = 0.676). I, same as C, but D_1_-like dopamine receptor antagonist SCH 23390 (100 µM) was bath-applied prior to pairing and continuously until the end of experiments (n = 5). Sample traces (right) include an example recorded after application of SCH 23390 but before pairing (at −2 min; blue). J, quantification of the data shown in I. Pairing was ineffective in the presence of SCH 23390, while SCH 23390 alone did not affect first EPSC amplitudes (p = 0.791, repeated measures one-way ANOVA) or PPRs (p = 0.464). K, same as I, but instead of SCH 23390, only the solvent DMSO (0.1%) was bath-applied (n = 6). L, quantification of the data shown in I. The effects of pairing were unaffected by DMSO, while DMSO alone did not affect first EPSC amplitudes or PPRs. P values for EPSCs are (from left to right) 0.973, < 10^−4^, < 10^−4^ and < 10^−4^ (Dunnett’s multiple comparisons test following repeated measures one-way ANOVA, p < 10^−6^), and for PPRs, 1.00, 0.00896, 0.00304 and < 10^−3^ (repeated measures one-way ANOVA, p < 10^−3^).

**Figure 3.**
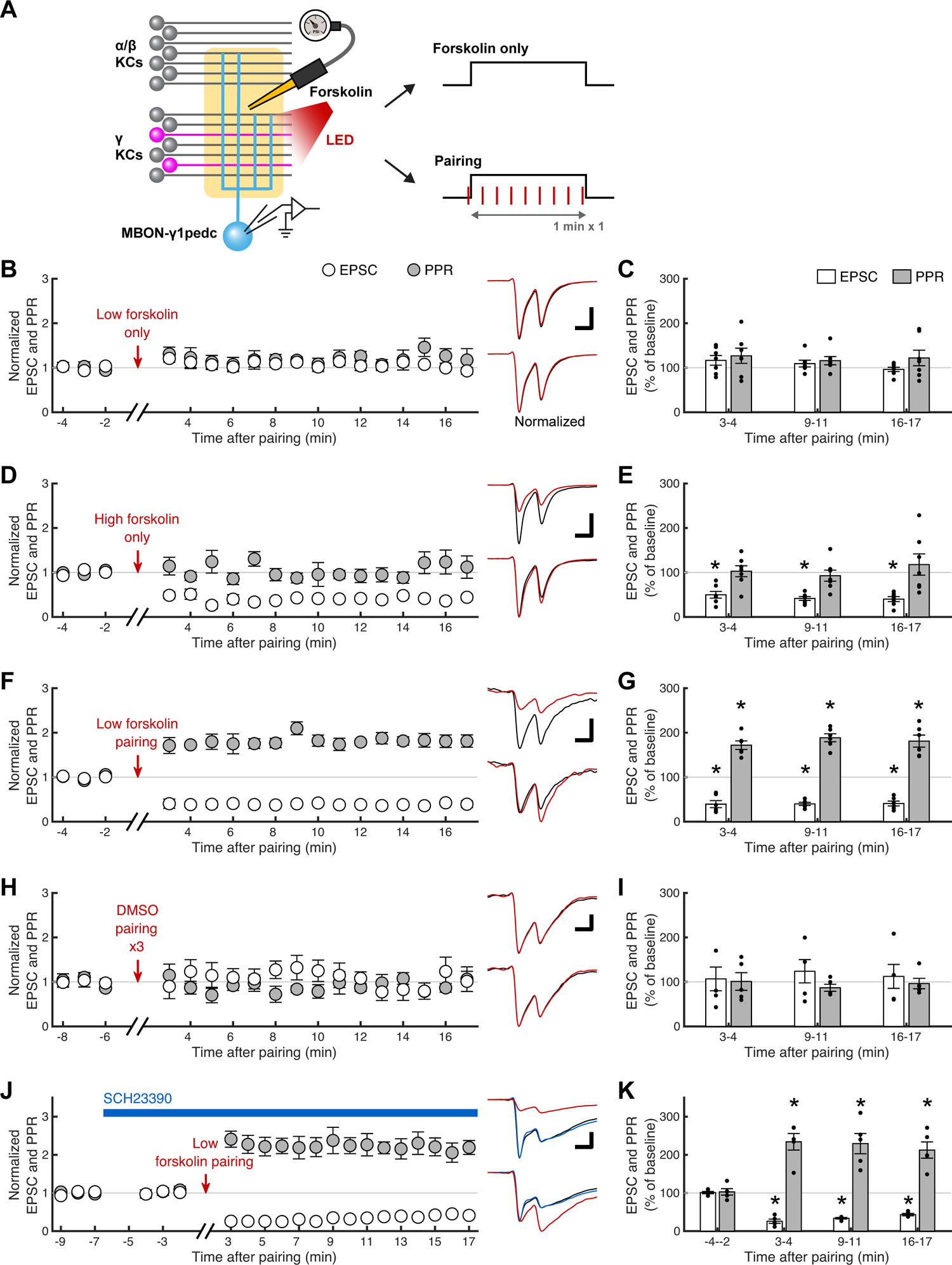
Presynaptic LTD induction at **γ** KC-to-MBON-**γ**1pedc synapses requires both AC activation and KC activity A, a schematic of the experiments. AC activator forskolin was focally applied to the dendritic region of the MBON-γ1pedc via an injection pipette while measuring optogenetically evoked γ KC-to-MBON-γ1pedc EPSCs. B, first EPSC amplitudes (open circles, mean ± SEM; n = 7) and PPRs (filled circles) plotted against time after the end of 1-min injection of forskolin (20 µM). The data were normalized to the average of a 3-min baseline recorded before pairing. Upper right traces show overlaid representative EPSCs sampled before (at −2 min; black) and after (at 3 min; red) injection. Horizontal and vertical scale bars in this and the other panels indicate 300 ms and 30 pA, respectively. Lower right traces show the same EPSCs normalized with the first EPSC amplitude. C, quantification of the data shown in B at early, middle and late periods after injection (mean ± SEM). Black dots indicate data from individual cells. 1-min injection of a low concentration of forskolin alone affected neither first EPSC amplitudes (open bars; p = 0.0866, repeated measures one-way ANOVA) nor PPRs (filled bars; p = 0.553). D, same as B, but with a higher concentration of forskolin (100 µM; n = 7). E, quantification of the data shown in D. 1-min injection of a high concentration of forskolin decreased first EPSC amplitudes at all three time points (p = 0.0151, 0.00645, and 0.00495 from left to right, Dunnett’s multiple comparisons test following repeated measures one-way ANOVA, p = 0.00502) but did not affect PPRs (p = 0.494, repeated measures one-way ANOVA). F, same as B, but 1-min forskolin (20 µM) injection was paired with γ KC activation (n = 6). G, quantification of the data shown in F. 1-min pairing of a low concentration of forskolin and γ KC activation depressed first EPSC amplitudes and increased PPRs at all three time points. P values for EPSCs are (from left to right) < 10^−6^, < 10^−6^ and < 10^−6^ (Dunnett’s multiple comparisons test following repeated measures one-way ANOVA, p < 10^−7^), and for PPRs, < 10^−3^, < 10^−4^ and 10^−3^ (repeated measures one-way ANOVA, p < 10^−4^). H, same as B, but instead of forskolin, only the solvent DMSO (0.1 %) was injected (n = 5). 1-min injection was repeated 3 times with 1-min intervals so that the data could also serve as control for the experiments shown in Fig. 7. I, quantification of the data shown in H. DMSO injection alone did not affect first EPSC amplitudes (p = 0.908, repeated measures one-way ANOVA) or PPRs (p = 0.708). J, same as F, but SCH 23390 (100 µM) was bath-applied prior to pairing and continuously until the end of experiments (n = 5). Sample traces (right) include an example recorded after application of SCH 23390 but before pairing (at −2 min; blue). K, quantification of the data shown in J. The effects of pairing were unaffected by SCH 23390, while SCH 23390 alone did not affect first EPSC amplitudes or PPRs. P values for EPSCs are (from left to right) 1.000, < 10^−8^, < 10^−7^ and < 10^−6^ (Dunnett’s multiple comparisons test following repeated measures one-way ANOVA, p < 10^−9^), and for PPRs, 1.00, <10^−4^, < 10^−4^ and < 10^−4^ (repeated measures one-way ANOVA, p < 10^−5^).

### Electrophysiology

We first attempted to perform all experiments *in vivo*. However, light stimulation we used for optogenetic activation of KCs evoked an EPSC-like inward current in MBON-γ1pedc as well as many of the randomly selected neurons in flies without CsChrimson transgene (data not shown). These responses persisted even in the presence of TTX (1 µM), but they almost disappeared in blind *norpA* mutants and were completely absent when we removed the retina. We therefore decided to switch to the *ex vivo* preparation. This strategy also improved the recording condition by minimizing the spontaneous circuit activity.

We dissected out a brain from a head capsule in ice-cold external saline, which contains (in mM) 103 NaCl, 3 KCl, 1.5 CaCl_2_, 4 MgCl_2_, 26 NaHCO_3_, 5 TES, 1 NaH_2_PO_4_, 10 trehalose and 10 glucose (pH 7.3 when bubbled with 95% O_2_ and 5% CO_2_, 275 mOsm), and then transferred it to a recording chamber, where the brain was pinned to the Sylgard-coated bottom using sharpened tungsten rods inserted at the optic lobes. In some experiments, we treated the brain with Type IV collagenase (0.2-0.5 mg/ml) for 30-90 s to make it easier to remove the glial sheath. After removing the sheath around the region of interest by forceps under a dissection scope or by pipettes under an upright microscope (OpenStand; Prior Scientific) equipped with a 60× water-immersion objective (LUMPlanFl/IR; Olympus), we inserted an injection pipette containing external saline with Texas Red-conjugated dextran (3,000 MW; 100-200 µM) and additional drugs as described in each figure legend. The tip of the injection pipette was placed near the dendritic region of MBON-γ1pedc under the guidance of the GFP signal in the MBON. Both injection and patch pipettes were made from borosilicate glass capillaries with filament (Sutter Instrument) by a micropipette puller (P-97, Sutter Instrument). Patch pipettes were further heat-polished at the tip and had a resistance of 3.5-5.5 MΩ. Pipettes with a slightly larger tip diameter were used as injection pipettes. Patch pipettes were filled with internal saline containing (in mM) 140 cesium aspartate, 10 HEPES, 1 EGTA, 1 KCl, 4 Mg-ATP, 0.5 Na-GTP and 10 QX-314 (pH adjusted to 7.3 with CsOH, 265 mOsm). Whole-cell voltage-clamp recordings were made from MBON-γ1pedc using Axon MultiClamp 700B amplifier (Molecular Probes). Cells were held at −60 mV. Leak current was typically < 150 pA. Series resistance was compensated up to 70% so that the uncompensated component remains constant at around 5 MΩ. Signals were low-pass filtered at 5 kHz before being digitized at 10 kHz. Sample traces shown in figures were further low-pass filtered.

KCs were stimulated by 1-ms light pulses delivered through the objective at 4.25 mW/mm^2^ by a high-power LED source (LED4D067; Thorlabs) equipped with 625 nm LED. To measure PPR, we first recorded a single-pulse EPSC to be used as a reference trace. After 10 s, we delivered 4 paired pulses with 400-ms intervals every 10 s. We repeated this set every minute. A second EPSC waveform was obtained every minute by subtracting a reference trace from the average of 4 paired-pulse EPSCs. PPR was then calculated every minute by dividing the second EPSC amplitude by the first EPSC amplitude. To ensure the stability of response, we recorded baseline responses for at least 3 min. For focal injection of drugs, we applied 1-s pressure pulses (0.4-0.6 psi) every 2 s for 1 min by a microinjector (PV850, World Precision Instruments). In some experiments, we repeated it three times with 1-min intervals. When we paired KC activation with focal injection, 1-ms photostimuli were delivered at 2 Hz for 1 min with the first pulse delivered 0.3 s before the onset of the first injection pulse. Recording was resumed 2.5 min after the end of injection. For bath application of drugs, we waited for 2 min after the normal bath saline was exchanged with the one containing a drug via perfusion. We then recorded EPSCs for 3 min to assess the effect of the drug itself before starting the pairing procedure described above. All stimulus delivery and data acquisition were controlled by custom MATLAB (Mathworks) codes. Data analyses were also performed on MATLAB. Statistical analyses were performed on MATLAB or Prism (GraphPad). The time course data of EPSCs and PPRs are shown in normalized values because the initial EPSC size was highly variable presumably because of our stochastic labeling strategy of KCs; initial EPSC size did not correlate with initial PPR (data not shown). All statistical tests used raw data before normalization.

### Two-photon imaging of cAMP

cAMP imaging was performed *in vivo* as described previously (Davidson *et al*., 2023) using a custom resonant-scanning two-photon microscope equipped with a femtosecond laser fixed at 930 nm (Menlo Systems) and a 20×, 1.00 NA water-immersion objective (XLUMPLFLN; Olympus). Optogenetic stimulation and forskolin or dopamine injection were done in the same way as the electrophysiological experiments except that Alexa 594 (50 µM) instead of Texas Red dextran was infused in the pipette solution to facilitate the visualization of the tip of the injection pipette under two-photon excitation. 512-by-512-pixel images were acquired at 30 frames per second and using ∼30 mW laser power. To protect photomultiplier tubes (PMTs), we closed the mechanical shutters installed in front of the PMTs during every photo-stimulation pulse.

Imaging data were preprocessed and analyzed by ImageJ (Schindelin *et al*., 2012) and custom MATLAB codes. To analyze the CsChrimson::tdTomato-positive and negative axons separately, we took the following steps. We first excluded the frames acquired during the injection pulses because the tissue was transiently deformed. We then averaged and median-filtered (3-by-3 pixels) red-channel images (CsChrimson::tdTomato) to use them as a reference to create CsChrimson-positive and negative binary masks, which were used to analyze the green-channel images (i.e. G-Flamp1). Since there was a small but steady drift of the tissue, we updated the mask every 6 s; during each 6-s period, the drift was negligible. We conservatively created CsChrimson-positive masks by selecting pixels with top ∼10% intensities after excluding the pixels of Alexa 594-positive pipette. CsChrimson-negative masks were from the bottom ∼50%, which was also conservative given half the pixels in this category were essentially empty. These masks were multiplied with the G-Flamp1-positive binary mask (top ∼50%), which matched the γ lobe structure, and used for analyses. Responses are presented as z-scores calculated based on a 3-s baseline period immediately before the stimulus period. A condition was judged as sufficient to evoke a cAMP response based on whether the mean response during the period between the stimulus onset and 15 s after the offset was significantly different from 0, as determined by a single-sample t-test for each condition (Forskolin + LED, LED Only, and Dopamine + LED). “LED Only” data were acquired from the same flies as the “Forskolin + LED”, following a 10-min waiting period. We also repeated the experiment with “LED Only” first and obtained equivalent results (n = 4; data not shown). Statistical analyses were performed on Prism.

### Drugs

Drug-containing external saline was freshly prepared on the day of experiment from stock solutions stored at −20 °C. Stock solutions of mecamylamine, Rp-cAMPS and KN-93 phosphate were made with water at 100 mM, and SCH 23390, forskolin, H-89 and BAY 41-2272 were dissolved in DMSO at 100 mM. Final concentration of DMSO did not exceed 0.1%. Dopamine was stored at 100 mM in external saline. When a drug was bath applied, that drug was also included in the injection pipette at the same concentration.

## Results

### Optogenetic paired-pulse stimulation of KCs can be used to assess presynaptic changes in synaptic transmission

In the field of synaptic physiology, paired-pulse protocol is commonly used to assess the presynaptic strength of synaptic transmission because the paired pulse ratio (PPR), calculated as the second EPSC amplitude divided by the first one, generally inversely correlates with presynaptic vesicular release probability (Zucker & Regehr, 2002). Although paired EPSCs are typically evoked by fiber stimulation using extracellular electrodes, similar inference can be made using optogenetically delivered paired pulses (Britt *et al*., 2012; Creed *et al*., 2016; Liu *et al*., 2020). This approach is applicable to *Drosophila* KCs whose densely packed, small bundle of axons deters the use of an electrode. Since this method has never been used in the *Drosophila* MB, to our knowledge, we first asked whether release probability change can induce predicted change in the PPR.

Among dozens of MBONs, we targeted MBON-γ1pedc because the relatively thick and short primary neurite of this neuron allows for superior membrane voltage control (i.e. space clamp) during somatic voltage-clamp recordings and also because the LTD has been best characterized in this MBON using pairing of odor and DAN activation (Hige *et al*., 2015). MBON-γ1pedc receives the majority of its inputs from the γ subtype of KCs as well as a minor fraction from α/β KCs in the pedunculus region of the MB (Fig. 1A). We therefore first focused on γ KC-to-MBON-γ1pedc synaptic transmission. To selectively study these synapses, we expressed red-shifted channelrhodopsin, CsChrimson (Klapoetke *et al*., 2014), in a small subset of γ KCs using γ KC-specific split-GAL4 driver MB623C (Shuai *et al*., 2023) together with a stochastic expression system SPARC (Isaacman-Beck *et al*., 2020). By using the “S” (or sparse) variant of the SPARC system, we can reliably label a random ∼3-7% of γ KCs (Isaacman-Beck *et al*., 2020), roughly equivalent to the fraction of KCs reliably responsive to a typical odor (Honegger *et al*., 2011).

We made whole-cell voltage-clamp recordings from MBON-γ1pedc and delivered two 1-ms light pulses 400 ms apart to measure PPRs. To test the effect of release probability decrease on PPR, we changed the extracellular calcium/magnesium concentrations from 1.5/4 to 0.7/5.5 mM. This manipulation decreased the first EPSC amplitude while concurrently increasing the PPR (Fig. 1B). Conversely, increasing the release probability by changing calcium/magnesium concentrations to 5/0.5 mM facilitated the first EPSC while decreasing the PPR (Fig. 1C). Thus, artificial manipulation of release probability shifted PPR in the expected manner. To test whether the change in PPR is attributable to presynaptic factors, we next bath-applied a low concentration of mecamylamine (10 µM), a non-competitive antagonist of the nicotinic acetylcholine receptors, to reduce the availability of postsynaptic ionotropic receptors without changing the release probability. This manipulation attenuated the first EPSC to an equivalent level to the low calcium condition (Fig. 1D). However, this decrease was not accompanied by a change in PPR. These results indicate that PPRs measured by optogenetically evoked EPSCs at KC-to-MBON synapses can be used as an indicator of presynaptic modulation of transmission.

### Pairing **γ** KC activation with focal dopamine application induces presynaptic LTD

Using this experimental setup, we first examined whether dopamine can induce synaptic plasticity at γ KC-to-MBON-γ1pedc synapses. Previous studies that demonstrated odor-specific depression in MBON responses used either actual reinforcement or direct DAN activation using opto- or chemogenetics (Séjourné *et al*., 2011; Owald *et al*., 2015; Hige *et al*., 2015; Cohn *et al*., 2015; Perisse *et al*., 2016; Berry *et al*., 2018; Felsenberg *et al*., 2018; Handler *et al*., 2019; Awata *et al*., 2019; Zhang *et al*., 2019; McCurdy *et al*., 2021; Hancock *et al*., 2022; Schnitzer *et al*., 2022; Noyes & Davis, 2023; Zeng *et al*., 2023), which can promote release of not only dopamine but also cotransmitters (Aso *et al*., 2019) or other neuromodulators. We therefore do not know whether dopamine alone is sufficient for the DAN activation-induced LTD. To directly test this possibility, we focally applied dopamine (1 mM) into γ1pedc subregion of the MB lobe, where the dendrites of MBON-γ1pedc are located, by pressure injection via a pipette placed in the MB lobe (Fig. 2A). By monitoring the signal of Texas Red-conjugated dextran infused in the pipette, we confirmed that our 1-min injection protocol (30 cycles of 1-s on and 1-s off) is enough to diffuse the injected solution across the entire γ1pedc, but it was largely confined to half the length of the medial MB lobes (Fig. 2B). Importantly, the signal was quickly washed out before resuming EPSC recording 2.5 min after the end of injection. After we paired optogenetic activation of γ KCs with dopamine application for 1 min, the synapses underwent LTD (54.7 ± 8.5 % depression after 3-4 min, mean ± SEM; n = 6), which lasted at least for 17 min (Figs. 2C and 2D). In 2 of the 6 recordings, we were able to continue the recording for more than 30 min after pairing. In both these cases, the LTD persisted until the end of recording without any sign of recovery (data not shown). This LTD was accompanied by PPR increase (91.4 ± 16.6 % increase), which persisted for the duration of LTD (Figs. 2C and 2D). In contrast, dopamine application (Figs. 2E and 2F) or γ KC activation alone (Figs. 2G and 2H) induced no change in EPSC amplitude or PPR. These results indicate that coincidence of γ KC activation and dopamine input can induce presynaptic LTD. To test if the action of dopamine is mediated by G_s_-coupled D_1_-like dopamine receptors, we bath-applied a selective antagonist SCH 23390 (100 µM). Although application of SCH 23390 by itself had no effect on the EPSC amplitude or PPR, it abolished the effect of γ KC-dopamine pairing (Figs. 2I and 2J). In contrast, bath application of the solvent DMSO (0.1%) did not have such effects, as γ KC-dopamine pairing still induced robust LTD and PPR increase (Figs. 2K and 2L). Taken together, these results indicate that the role of DAN activation in LTD induction described in previous studies (Hige *et al*., 2015; Cohn *et al*., 2015; Berry *et al*., 2018; Handler *et al*., 2019) is attributable to dopamine’s action on D_1_-like dopamine receptors in these synapses.

### Presynaptic LTD induction requires both AC activation and KC activity

The dominant hypothesis in the field is that coincidence of KC activity and dopamine input activates AC in KC axons to elevate cAMP concentration which in turn induces synaptic plasticity. This model assumes that cAMP increase is sufficient for plasticity induction. To test this long-standing but unproved hypothesis, we pharmacologically activated the AC by focally injecting forskolin (20 µM) into γ1pedc region (Fig 3) as we did for dopamine. As we will show later, forskolin at this concentration elicited the reliable cAMP response in KC axons. However, 1-min injection of forskolin affected neither the EPSC amplitude nor PPR (Figs. 3B and 3C).

Increasing the concentration of forskolin to 100 µM did induce a sustained decrease in the EPSC amplitude (Figs. 3D and 3E). However, this effect did not accompany a change in PPR, suggesting postsynaptic origin of the plasticity. It is possible that excessive concentration of forskolin recruited AC in the MBON. In support of this idea, cell-type-specific transcriptome data (Aso *et al*., 2019) suggest that expression level of AC is much higher in KCs compared to the MBON (Fig. 4). Although the lack of change in PPR does not formally exclude the possibility that high concentrations of forskolin induces plasticity through presynaptic mechanisms, we can at least conclude that the observed plasticity is qualitatively distinct from the plasticity induced by γ KC-dopamine pairing. In other words, elevation of cAMP caused by AC activation is not sufficient to replicate the dopamine-induced LTD.

**Figure 4.**
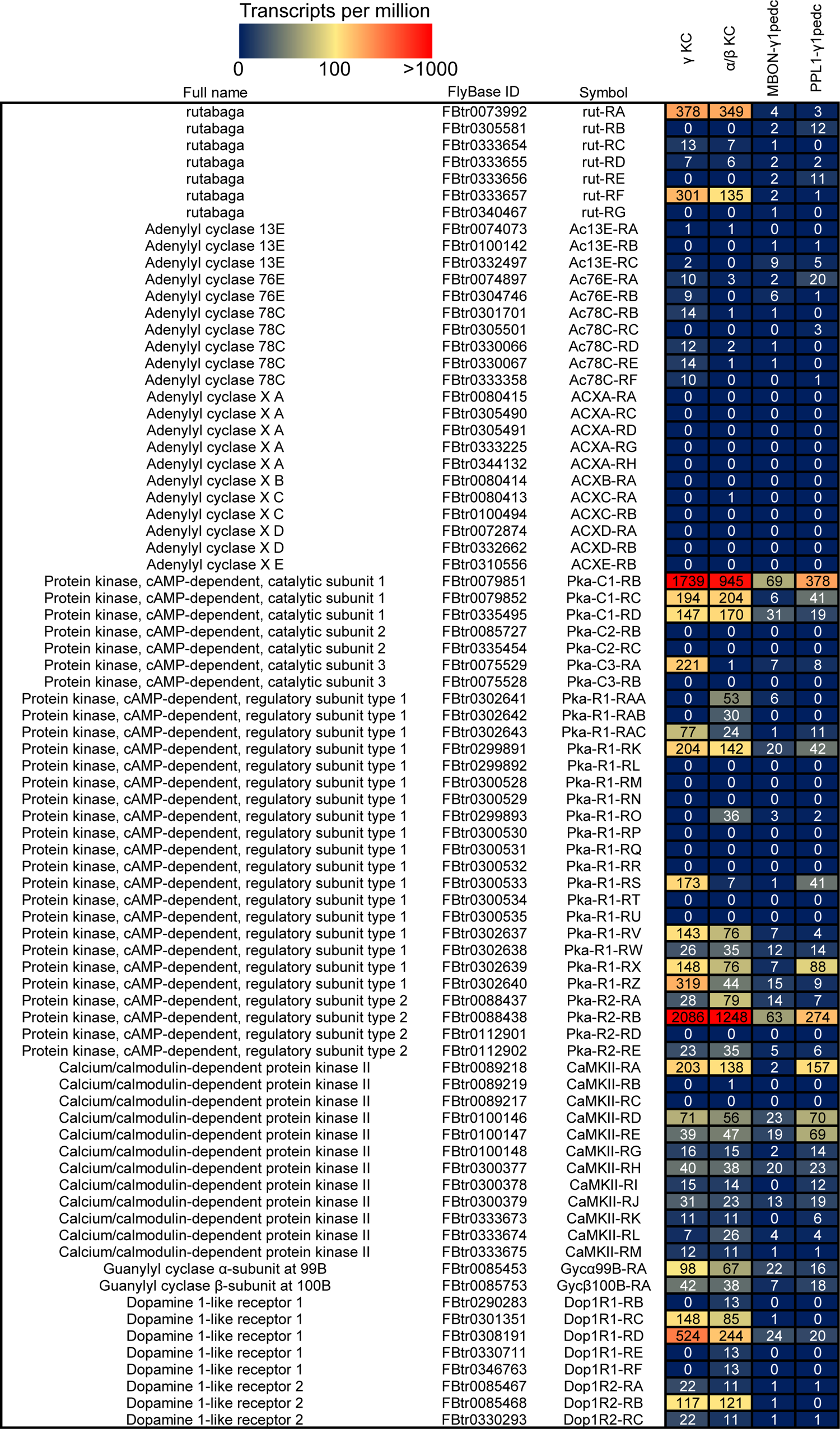
Transcriptome data related to pharmacological target molecules Cell-type-specific transcriptome data of the genes encoding the target molecules of pharmacology used in this study. This figure was recreated based on published data (Aso *et al*., 2019). PPL-γ1pedc is the DAN whose axonal innervation pattern in the MB lobes matches the dendritic arborization of MBON-γ1pedc.

These unexpected results prompted us to test whether KC activation in addition to cAMP elevation is necessary to induce presynaptic LTD. To this end, we paired γ KC activation and injection of a low concentration of forskolin. This pairing was able to induce a long-lasting suppression (at least for 17 min) of the EPSC amplitude (60.6 ± 8.0 % depression after 3-4 min; n = 6) and concurrent increase in PPR (71.9 ± 9.6 % increase; Figs. 3F and 3G), replicating dopamine-induced LTD. In contrast, pairing of γ KC activation with injection of DMSO (0.1%; solvent of forskolin) did not show any effect (Figs. 3H and 3I). It is possible that during KC-forskolin pairing, forskolin facilitates dopamine release from DANs, which would reproduce practically the same condition as KC-dopamine pairing. To test this possibility, we bath-applied SCH 23390 (Figs. 3J and 3K). Unlike the case with KC-dopamine pairing (Figs. 2I and 2J), SCH 23390 did not affect the LTD induced by KC-forskolin pairing. Thus, the KC-forskolin pairing protocol does bypass the activation of dopamine receptors to induce LTD. These results suggest that some intracellular signal triggered by KC activity is required to converge somewhere in the downstream pathway of the cAMP production to express presynaptic LTD.

### KC activation does not exert additive effect on forskolin- or dopamine-induced presynaptic cAMP elevation

Our conclusion above is based on the assumption that forskolin application occludes further activation of AC by neuronal activity. However, our results do not formally rule out the possibility that forskolin and KC activity exert a synergistic effect on cAMP level and thereby gate LTD. To test this possibility, we performed cAMP imaging. We expressed the cAMP sensor G-Flamp1 (Wang *et al*., 2022) in all γ KCs using MB623C and CsChrimson in a sparse subset of the γ KCs using the SPARC system. We then paired focal injection of forskolin (20 µM) into γ1pedc region and photo-activation of γ KCs using the same protocol as the electrophysiological experiments (Fig. 5A). To test the effect of KC activation, CsChrimson-negative and positive axons were analyzed separately (Figs. 5B and 5C; see also Methods). Pairing induced robust cAMP increase in both CsChrimson-negative and positive KC axons (Fig. 5D). The cAMP level largely returned to the baseline within 2-3 min after pairing. This decay time course is consistent with that of Texas Red signal infused in the injection pipette in electrophysiological experiments (Fig. 2B), suggesting that maintenance of presynaptic LTD does not require sustained cAMP elevation. Importantly, there was no difference in the response magnitude between CsChrimson-negative and positive axons (Fig. 5G), indicating that KC activation does not facilitate forskolin-induced cAMP synthesis. Furthermore, KC activation alone did not evoke any cAMP increase (Figs. 5E and 5G). These results strengthen our conclusion that simple cAMP increase cannot induce presynaptic LTD. Next, we tested whether dopamine and KC activity can synergistically increase presynaptic cAMP level, which has been reported in multiple studies (Tomchik & Davis, 2009; Gervasi *et al*., 2010; Handler *et al*., 2019). Pairing of dopamine (1 mM) and γ KC activation increased the G-Flamp1 signal in CsChrimson-negative KC axons (Fig. 5F), indicating that dopamine injection alone can induce robust cAMP increase without coincident neuronal activity. The pairing also elicited cAMP increase in CsChrimson-positive axons. The magnitude of this response was not different from the forskolin-induced response in CsChrimson-negative axons (Fig. 5G), which further strengthens the point that cAMP is not a sufficient trigger of LTD. Surprisingly, there was no difference in the response magnitude between CsChrimson-negative and positive axons (Fig. 5G), providing no evidence for synergistic action of dopamine input and neuronal activity on cAMP synthesis. These results are largely in agreement with the recent study that also demonstrated the lack of difference in the cAMP level between odor-responding KC axons and the other KCs during odor-shock pairing (Abe *et al*., 2023). Taken together, we conclude that the convergence of intracellular signaling triggered by dopamine input and neuronal activity, which represents crucial coincidence detection of reinforcement and olfactory signals during classical conditioning, takes place after, rather than to trigger, cAMP synthesis.

**Figure 5.**
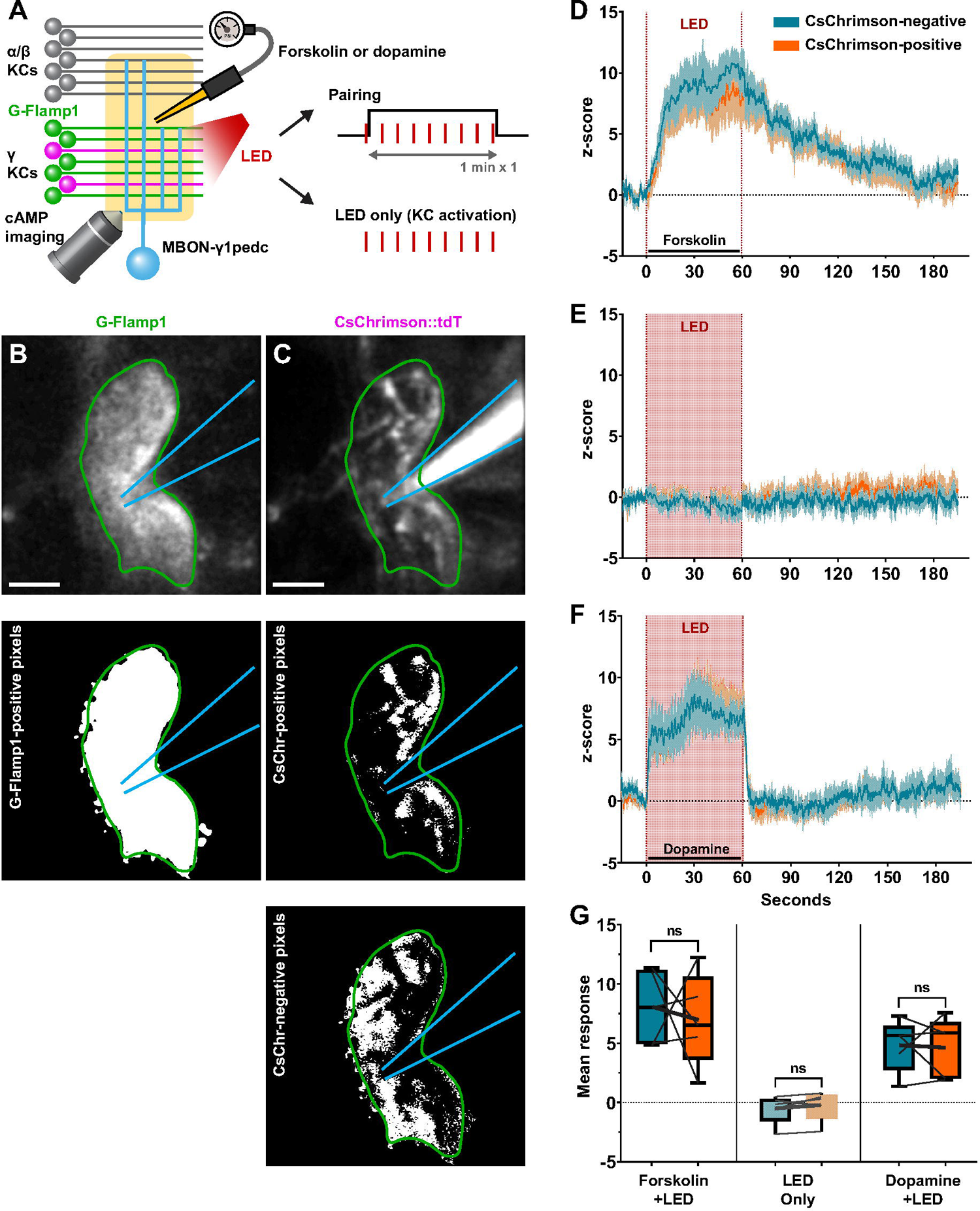
KC activation does not exert additive effect on forskolin- or dopamine-induced presynaptic cAMP elevation A, a schematic of the experiments. Forskolin (20 μM) or dopamine (1 mM) was paired with γ KC activation while measuring cAMP level via in vivo, two-photon imaging of the genetically encoded cAMP sensor, G-Flamp1. In a subset of experiments, the LED only condition was also tested. B, Upper, representative field of view in the green channel, which shows G-Flamp1 signal. Green outline denotes approximate boundaries of the γ1 region, and blue outline denotes approximate injection pipette boundaries. Lower, binary mask created from the same image to define the γ1 lobe. Scale bar, 5µm. C, Upper, same as B but in the red channel, which visualizes CsChrimson:tdTomato and Alexa 594 signal. Middle, binary mask to define CsChrimson-positive pixels. After manually excluding the pixels representing the Alexa 594-marked injection pipette, pixels with top 10% intensity were selected and then multiplied by the mask in B. Lower, binary mask to define CsChrimson-negative pixels. Those with bottom 50% pixel intensity were selected and multiplied by the mask in B. See Methods for the detail. D, Response profile of CsChrimson-negative (teal) and -positive (orange) axons in response to pairing of forskolin (20 μM) injection and KC activation (mean ± SEM; n = 5). E, Same as D for KC activation alone (n = 5). F, Same as D for pairing of dopamine (1 mM) injection and KC activation (n = 5). G, Comparison of mean response for each group of axons. Pairing with forskolin (Forskolin + LED; p < 10^−4^, single-sample t-test) and dopamine (Dopamine + LED; p < 10^−4^) evoked significant changes in cAMP levels in both CsChrimson-negative (teal) and -positive (orange) axons, while KC activation alone (LED Only; p = 0.344) did not evoke a response. The magnitude of response did not differ between CsChrimson-positive and -negative axons in any of the experiments (forskolin, p = 0.730, paired t-test; LED Only, p = 0.235; dopamine, p = 0.917). The magnitude of the forskolin-evoked response was not different from the dopamine-evoked response regardless of the combinations of CsChrimson expression (p = 0.0967 [Chrimson-negative forskolin *vs.* negative dopamine], 0.0989 [negative *vs.* positive], 0.313 [positive *vs.* negative] and 0.302 [positive *vs.* positive], unpaired t test).

### Dopamine-induced LTD depends on PKA

Our results so far demonstrated that the cAMP pathway is a critical molecular basis for LTD, even though its activation alone may not be sufficient. We therefore tested pharmacologically which molecules downstream of cAMP are required for LTD induction (Fig. 6A). As with many other organisms, PKA plays a crucial role in *Drosophila* olfactory learning (Drain *et al*., 1991; Skoulakis *et al*., 1993). However, its role in the LTD at the KC output synapse has not been examined. To test this, we first used Rp-cAMPS, a non-hydrolyzable cAMP analog that inhibits PKA activation. Bath application of Rp-cAMPS (100 µM) did not affect the EPSC amplitude or PPR but completely blocked the LTD induced by γ KC-dopamine pairing (Figs. 6B and 6C). We also tested the effect of another PKA inhibitor H-89. Bath-applied H-89 (10 µM) also abolished the LTD induced by γ KC-dopamine pairing without affecting baseline EPSC amplitude or PPR (Figs. 6D and 6E). As shown in Figs. 2K and 2L, bath application of the solvent of H-89 alone did not show any effect. Calcium/calmodulin-dependent protein kinase II (CaMKII) is another protein kinase that has a conserved role in synaptic plasticity across species (Bayer & Schulman, 2019). In *Drosophila*, it is also implicated in some form of associative learning other than olfactory learning (Griffith *et al*., 1993). Since our results suggest an important role of KC activity, which may lead to CaMKII activation via calcium influx, we tested the effect of a CaMKII inhibitor KN-93, which is effective in *Drosophila* (Peretz *et al*., 1998). However, in the presence of KN-93 (10 µM), γ KC-dopamine pairing induced robust, or even more pronounced, LTD and PPR increases (Figs. 6F and 6G). Since application of KN-93 itself slightly decreased the EPSC amplitude (but not PPR), phosphorylation by CaMKII may play a role in maintaining normal synaptic transmission, perhaps on the postsynaptic side. This suppression of the baseline EPSCs may partly explain why LTD is more pronounced. Taken together, dopamine-induced LTD in γ KCs depends on PKA, but CaMKII activation may not be critical for this form of synaptic plasticity.

**Figure 6.**
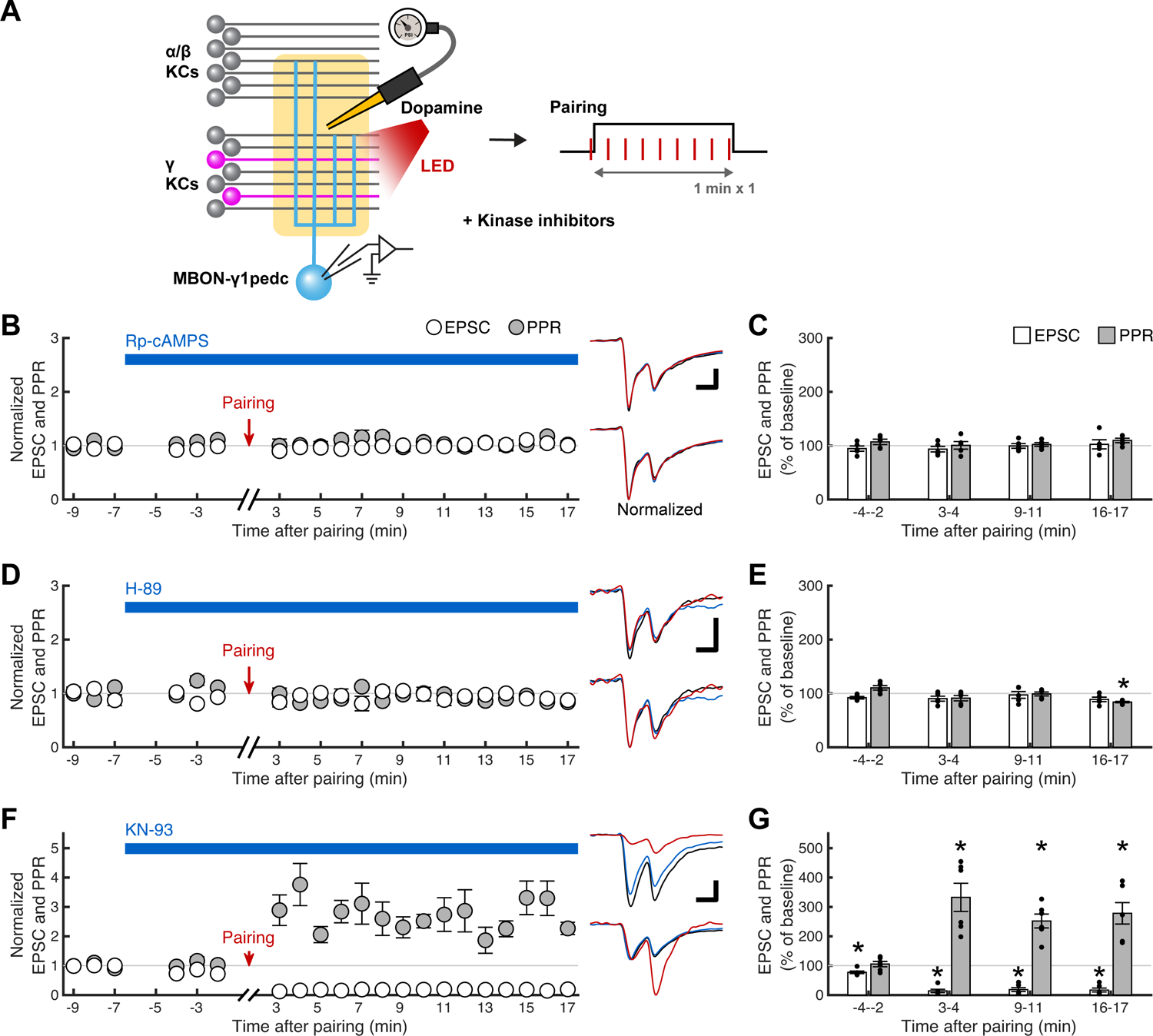
Dopamine-induced LTD depends on PKA but not CaMKII A, a schematic of the experiments. Dopamine (1 mM) injection was paired with γ KC activation while measuring optogenetically evoked γ KC-to-MBON-γ1pedc EPSCs as in Fig. 3C, except kinase inhibitors were bath-applied prior to pairing and applied continuously until the end of experiments. B, effect of a competitive antagonist of PKA, Rp-cAMPS (100 µM). First EPSC amplitudes (open circles, mean ± SEM; n = 5) and PPRs (filled circles) plotted against time after the end of 1-min pairing of γ KC activation and dopamine injection. The data were normalized to the average of a 3-min baseline recorded before pairing. A horizontal blue bar indicates the period of Rp-cAMPS application. Upper right traces show overlaid representative EPSCs sampled before (at −7 min; black) and 4.5 min after drug application (at −2 min; blue), and after pairing (at 3 min; red). Horizontal and vertical scale bars in this and the other panels indicate 300 ms and 30 pA, respectively. Lower right traces show the same EPSCs normalized with the first EPSC amplitude. C, quantification of the data shown in B before pairing and at early, middle and late periods after pairing (mean ± SEM). Black dots indicate data from individual cells. Rp-cAMPS alone did not affect first EPSC amplitudes (open bars) or PPRs (filled bars), and the subsequent pairing did not affect EPSCs (p = 0.335, repeated measures one-way ANOVA) or PPRs (p = 0.417). D, same as B, but instead of Rp-cAMPS, PKA inhibitor H-89 (10 µM) was bath-applied (n = 5). E, quantification of the data shown in D. H-89 alone did not affect first EPSC amplitudes or PPRs, and the subsequent pairing did not depress EPSCs (p = 0.392, repeated measures one-way ANOVA) or increase PPRs (p = 0.205, 0.171, 1.00 and 0.0174 from left to right, Dunnett’s multiple comparisons test following repeated measures one-way ANOVA, p < 10^−3^). F, same as B, but instead of Rp-cAMPS, CaMKII inhibitor KN-93 (10 µM) was bath-applied (n = 6). G, quantification of the data shown in F. KN-93 alone slightly depressed EPSCs without affecting PPRs, and the subsequent pairing further induced robust depression of EPSCs and facilitation of PPRs. P values for EPSCs are (from left to right) 0.0206, < 10^−8^, < 10^−8^ and < 10^−8^ (Dunnett’s multiple comparisons test following repeated measures one-way ANOVA, p < 10^−9^), and for PPRs, 1.00, < 10^−5^, < 10^−3^ and < 10^−3^ (repeated measures one-way ANOVA, p < 10^−5^).

### Simultaneous activation of cGMP pathway and **γ** KCs induces presynaptic LTP

A behavioral study in dopamine-deficient flies (i.e. the mutant flies that cannot synthesize dopamine in neurons) identified NO as a cotransmitter of a subset of DANs, including the one projecting to γ1pedc (Aso *et al*., 2019). Since pairing of odor presentation and optogenetic activation of those DANs in dopamine-deficient flies induced memories with opposite valence to normal flies, it has been hypothesized that cGMP pathway downstream of NO induces synaptic potentiation and opposes dopamine-induced LTD. To test this hypothesis, we used sGC agonist BAY 41-2272 (Fig. 7A), which can activate the *Drosophila* sGC consisting of Gycα99B/Gycβ100B subunits (Morton *et al*., 2005) expressed in KCs (Fig. 4) (Aso *et al*., 2019). When 1-min focal injection of BAY 41-2272 (100 µM) was repeated three times, it slowly potentiated the γ KC-to-MBON-γ1pedc synaptic transmission over ∼15 min in some cells, but this effect was not highly consistent between cells (Figs. 7B and 7C). This variable potentiation was not accompanied by a change in PPR. In contrast, when the same BAY 41-2272 injection pattern was paired with γ KC activation, we reproducibly observed slowly developing LTP (77.9 ± 11.9 % potentiation after 16-17 min; n = 5) with concurrent decrease in PPR (24.8 ± 5.3 % decrease; Figs. 7D and 7E). Thus, as reminiscent of the role of cAMP pathway in LTD, it requires simultaneous KC stimulation for activation of cGMP pathway to induce presynaptic plasticity, but the direction of plasticity is opposite to the one induced by cAMP. Another difference from cAMP-induced LTD is that one-time 1-min pairing of BAY 41-2272 injection with γ KC activation did not induce LTP (Figs. 7F and 7G). The requirement of multiple rounds of pairing for LTP induction and the slow kinetics of LTP are consistent with the behavioral study that showed that NO-dependent learning requires longer training than dopamine-dependent learning and that NO-dependent memory develops slowly over time, taking ∼10 min after training (Aso *et al*., 2019).

**Figure 7.**
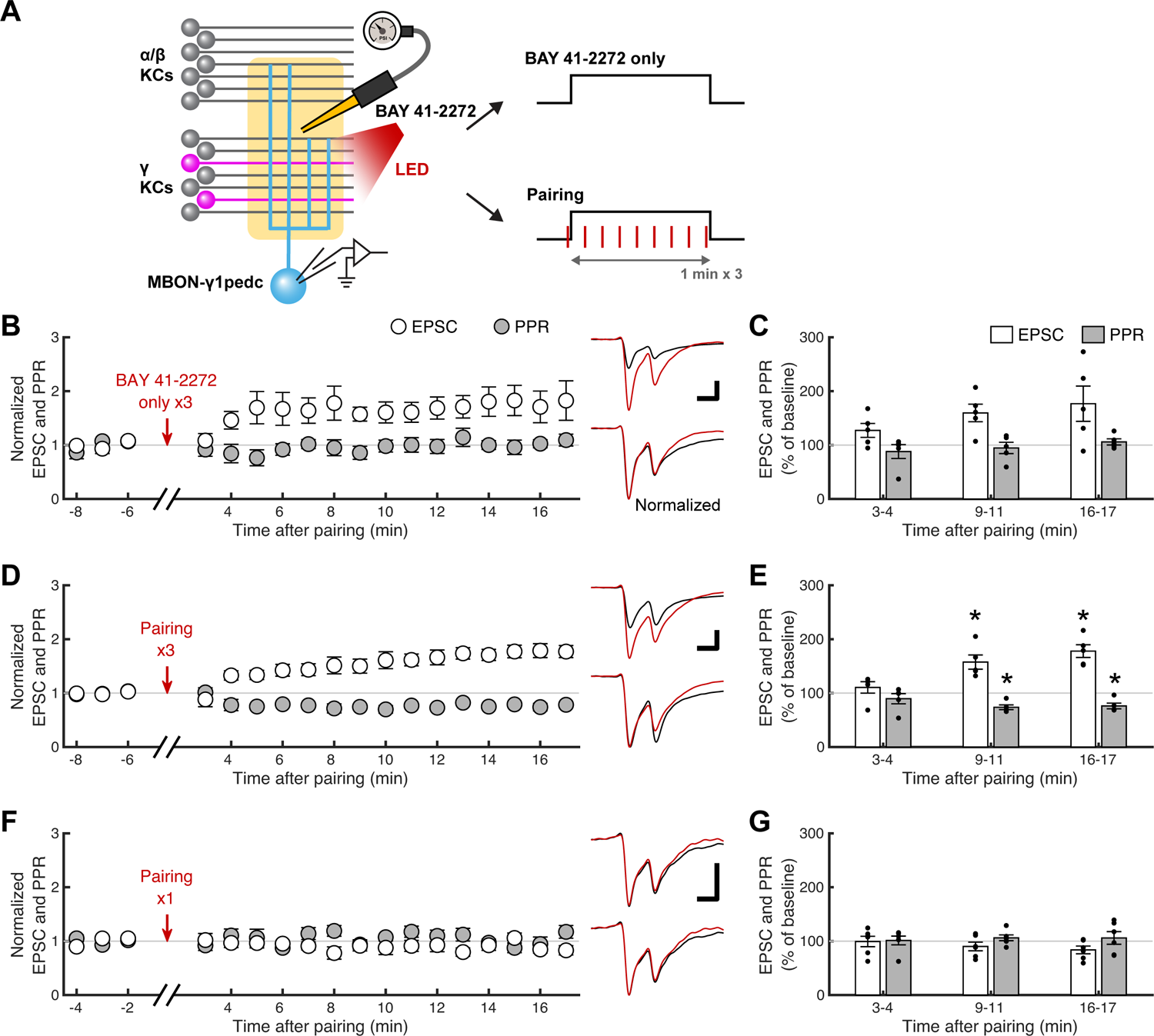
Simultaneous activation of cGMP pathway and **γ** KCs induces presynaptic LTP A, a schematic of the experiments. sGC agonist BAY 41-2272 was focally applied to the dendritic region of the MBON-γ1pedc via an injection pipette while measuring optogenetically evoked γ KC-to-MBON-γ1pedc EPSCs. B, first EPSC amplitudes (open circles, mean ± SEM; n = 5) and PPRs (filled circles) plotted against time after the end of the three repeats of 1-min injection of BAY 41-2272 (100 µM). The data were normalized to the average of a 3-min baseline recorded before pairing. Upper right traces show overlaid representative EPSCs sampled before (at −6 min; black) and after (at 17 min; red) injection. Horizontal and vertical scale bars in this and the other panels indicate 300 ms and 60 pA, respectively. Lower right traces show the same EPSCs normalized with the first EPSC amplitude. C, quantification of the data shown in B at early, middle and late periods after injection (mean ± SEM). Black dots indicate data from individual cells. BAY 41-2272 injection alone did not induce consistent changes in first EPSC amplitudes (open bars; p = 0.0633, repeated measures one-way ANOVA) or PPRs (filled bars; p = 0.565). D, same as B, but BAY 41-2272 injection was paired with γ KC activation (n = 5). E, quantification of the data shown in D. Pairing of BAY 41-2272 and γ KC activation potentiated first EPSC amplitudes and decreased PPRs at middle and late time points. P values for EPSCs are (from left to right) 0.314, < 10^−3^ and < 10^−3^ (Dunnett’s multiple comparisons test following repeated measures one-way ANOVA, p < 10^−4^), and for PPRs, 0.350, 0.00486 and 0.00902 (repeated measures one-way ANOVA, p = 0.00553). F, same as D, but pairing was performed only once (n = 6). G, quantification of the data shown in F. One-time pairing of BAY 41-2272 and γ KC activation was not sufficient to affect first EPSC amplitudes (p = 0.0932, repeated measures one-way ANOVA) or PPRs (p = 0.843).

### Depression at **α**/**β** KC-to-MBON-**γ**1pedc synapses is short lasting

In our plasticity induction method, odor-evoked KC activation, which occurs across all KC subtypes, is substituted with subtype-specific optogenetic activation. This feature allowed us to compare the properties of synaptic plasticity between different subtypes of KCs that share the same postsynaptic MBON. To test if the α/β KC-to-MBON-γ1pedc synapses undergo similar long-term synaptic plasticity to γ KC-to-MBON-γ1pedc synapses, we next expressed CsChrimson in a subset of α/β KCs using α/β KC-specific split-GAL4 driver MB008C (Aso *et al*., 2014b) and the SPARC system (Fig. 8A). As with the case of γ KCs, activation of α/β KCs paired with focal dopamine injection caused LTD (81.8 ± 3.8 % depression after 3-4 min; n = 6) with a concurrent increase in PPR (95.1 ± 11.1 % increase; Figs. 8B and 8C). However, the duration of LTD was markedly shorter. The EPSC amplitude as well as PPR started showing recovery within 5 min after induction, and both became indistinguishable from the baseline after ∼10 min. In contrast, at γ KC-to-MBON-γ1pedc synapses, LTD lasted at least for 30 min. Dopamine injection (Figs. 8D and 8E) or α/β KC activation (Figs. 8F and 8G) alone did not induce any plasticity. These results indicate that the properties of dopamine-induced synaptic plasticity are different between KC subtypes even among the synapses on the same MBON.

**Figure 8.**
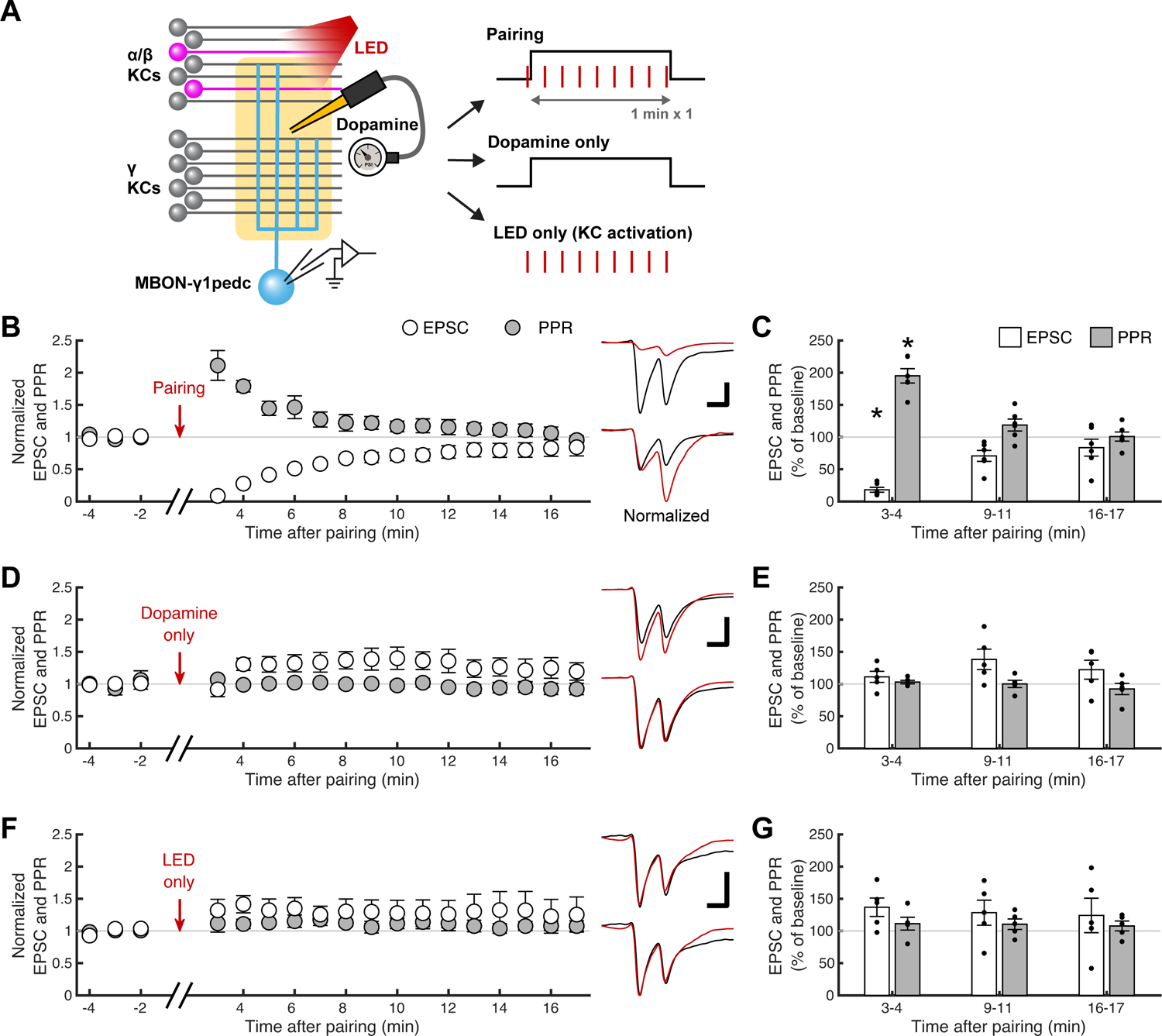
Pairing **α**/**β** KC activation with focal dopamine application induces transient presynaptic LTD A, a schematic of the experiments. Dopamine (1 mM) was focally applied to the dendritic region of the MBON-γ1pedc via an injection pipette while measuring optogenetically evoked α/β KC-to-MBON-γ1pedc EPSCs. B, first EPSC amplitudes (open circles, mean ± SEM; n = 6) and PPRs (filled circles) plotted against time after the end of 1-min pairing of α/β KC activation and dopamine injection. The data were normalized to the average of a 3-min baseline recorded before pairing. Upper right traces show overlaid representative EPSCs sampled before (at −2 min; black) and after (at 3 min; red) pairing. Horizontal and vertical scale bars in this and the other panels indicate 300 ms and 100 pA, respectively. Lower right traces show the same EPSCs normalized with the first EPSC amplitude. C, quantification of the data shown in B at early, middle and late periods after pairing (mean ± SEM). Black dots indicate data from individual cells. First EPSC amplitudes (open bars) and PPRs (filled bars) showed depression and increase, respectively, but only transiently at the early time point. P values for EPSCs are (from left to right) < 10^−4^, 0.100 and 0.593 (Dunnett’s multiple comparisons test following repeated measures one-way ANOVA, p < 10^−4^), and for PPRs, < 10^−5^, 0.253 and 0.999 (repeated measures one-way ANOVA, p < 10^−5^). D, same as B, but KC activation was omitted during pairing (n = 5). E, quantification of the data shown in D. 1-min dopamine injection alone affected neither first EPSC amplitudes (p = 0.208, repeated measures one-way ANOVA) nor PPRs (p = 0.350). F, same as B, but dopamine application was omitted during pairing (n = 5). G, quantification of the data shown in G. 1-min KC activation alone affected neither first EPSC amplitudes (p = 0.213, repeated measures one-way ANOVA) nor PPRs (p = 0.518).

To test whether difference in the duration of plasticity also applies to LTD induced by direct activation of AC, we next injected forskolin (Fig. 9A). Neither a low (10 µM; Figs. 9B and 9C) nor a high (100 µM; Figs. 9D and 9E) concentration of forskolin induced robust LTD or parallel increase in PPRs. In contrast, when we paired injection of a low concentration of forskolin with activation of α/β KCs, we observed transient but robust LTD (76.2 ± 9.5 % depression after 3-4 min; n = 5) accompanying parallel increase in PPRs (28.6 ± 10.0 % increase; Figs. 9F and 9G). Thus, as with the case of γ KCs, elevation of cAMP level by AC activation is not sufficient to induce LTD at α/β KC-to-MBON-γ1pedc synapses, as it additionally requires KC activation. On the other hand, the duration of LTD induced by α/β KC-forskolin pairing was reminiscent of that induced by α/β KC-dopamine pairing. To exclude the possibility that the short duration of LTD reflects insufficient diffusion of forskolin into the pedunculus subregion, where α/β KC-to-MBON-γ1pedc synapses are located, we repeated the same pairing experiment with a high concentration of forskolin (100 µM). The results mirrored those with the lower concentration; LTD and accompanying increase in PPRs were robust but still transient (Figs. 9H and 9I). Pairing of α/β KC activation with injection of DMSO (0.1%) did not show any effect (Figs. 9J and 9K). These results confirm the short-lasting nature of cAMP-induced LTD at α/β KC-to-MBON-γ1pedc synapses.

**Figure 9.**
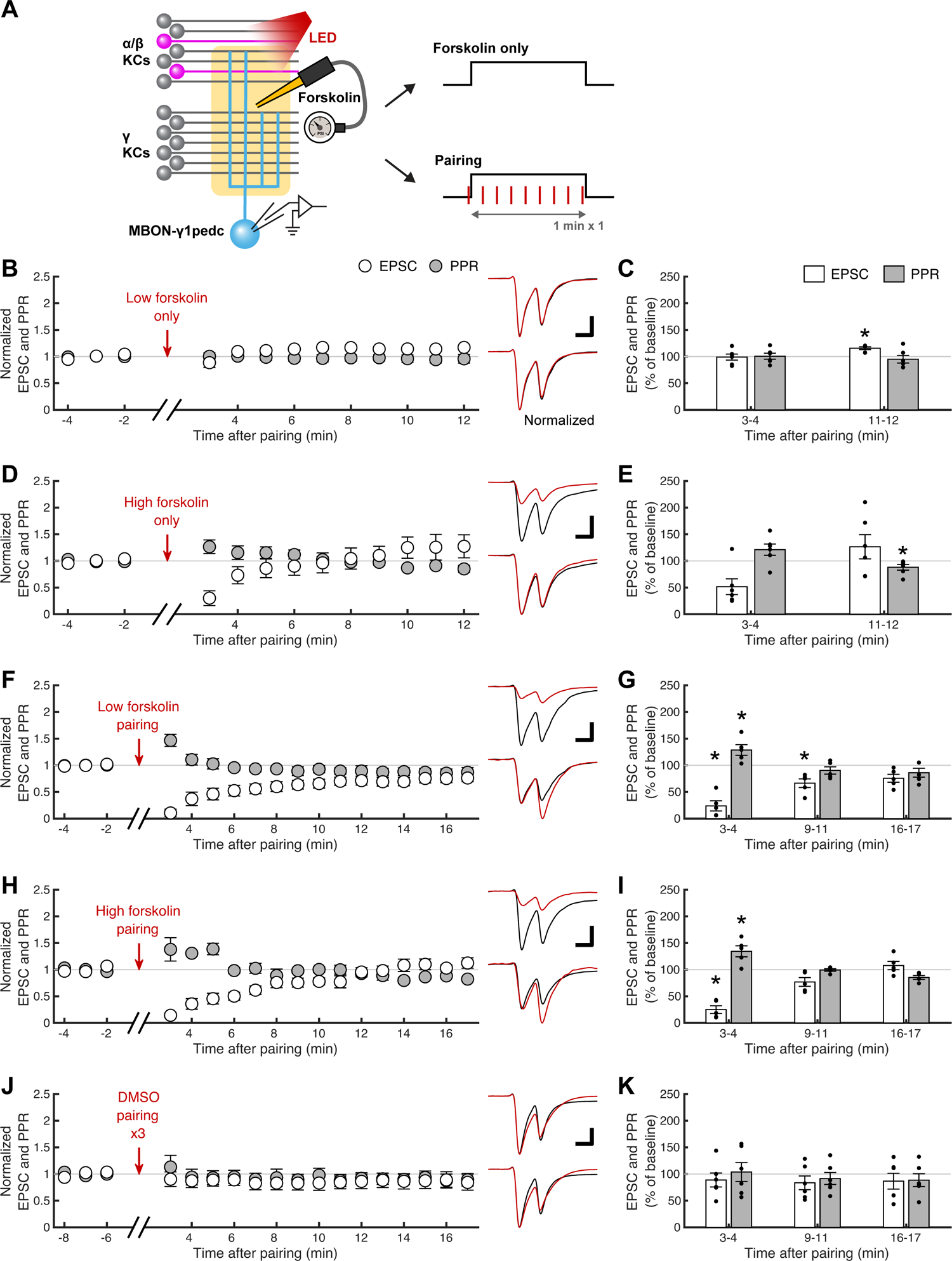
Presynaptic LTD induction at **α**/**β** KC-to-MBON-**γ**1pedc synapses requires both AC activation and KC activity A, a schematic of the experiments. Forskolin was focally applied to the dendritic region of the MBON-γ1pedc via an injection pipette while measuring optogenetically evoked α/β KC-to-MBON-γ1pedc EPSCs. B, first EPSC amplitudes (open circles, mean ± SEM; n = 6) and PPRs (filled circles) plotted against time after the end of 1-min injection of forskolin (10 µM). The data were normalized to the average of a 3-min baseline recorded before pairing. Upper right traces show overlaid representative EPSCs sampled before (at −2 min; black) and after (at 3 min; red) injection. Horizontal and vertical scale bars indicate 300 ms and 100 pA, respectively. Lower right traces show the same EPSCs normalized with the first EPSC amplitude. C, quantification of the data shown in B at early and late periods after injection (mean ± SEM). Black dots indicate data from individual cells. 1-min injection of a low concentration of forskolin alone did not induce consistent changes in first EPSC amplitudes (open bars; p = 0.999 and 0.0119 from left to right, Dunnett’s multiple comparisons test following repeated measures one-way ANOVA, p = 0.00922) and PPRs (filled bars; p = 0.512, repeated measures one-way ANOVA, p = 0.00922). D, same as B, but with a higher concentration of forskolin (100 µM; n = 6). Horizontal and vertical scale bars indicate 300 ms and 50 pA, respectively. E, quantification of the data shown in D. 1-min injection of a high concentration of forskolin alone did not induce consistent changes in first EPSC amplitudes (p = 0.0886, repeated measures one-way ANOVA) and PPRs (p = 0.0370 and 0.419 from left to right, Dunnett’s multiple comparisons test following repeated measures one-way ANOVA, p = 0.00796). F, same as B, but 1-min forskolin (10 µM) injection was paired with α/β KC activation (n = 5). Horizontal and vertical scale bars indicate 300 ms and 100 pA, respectively. G, quantification of the data shown in F. 1-min pairing of a low concentration of forskolin and α/β KC activation induced coherent but transient depression in first EPSC amplitudes and facilitation of PPRs. P values for EPSCs are (from left to right) < 10^−5^, 0.0105 and 0.0505 (Dunnett’s multiple comparisons test following repeated measures one-way ANOVA, p < 10^−4^) and for PPRs, 0.0117, 0.415 and 0.187 (repeated measures one-way ANOVA, p < 10^−3^). H, same as F, but with a higher concentration of forskolin (100 µM; n =5). Horizontal and vertical scale bars indicate 300 ms and 50 pA, respectively. I, quantification of the data shown in H. 1-min pairing of a high concentration of forskolin and α/β KC activation induced coherent but transient depression in first EPSC amplitudes and facilitation of PPRs. P values for EPSCs are (from left to right) < 10^−3^, 0.208 and 0.997 (Dunnett’s multiple comparisons test following repeated measures one-way ANOVA, p < 10^−3^) and for PPRs, 0.00258, 0.998 and 0.206 (repeated measures one-way ANOVA, p < 10^−3^). J, same as B, but instead of forskolin, only the solvent DMSO (0.1 %) was injected (n = 6). 1-min injection was repeated 3 times with 1-min intervals so that the data could also serve as control for the experiments shown in Fig. 10. Horizontal and vertical scale bars indicate 300 ms and 100 pA, respectively. K, quantification of the data shown in J. DMSO injection alone did not affect first EPSC amplitudes (p = 0.776, repeated measures one-way ANOVA) or PPRs (p = 0.535).

In contrast, cGMP-induced plasticity appeared similar between KC subtypes (Fig. 10A). As observed at γ KC synapses, three times 1-min injection of BAY 41-2272 alone did not induce LTP paralleled by PPR change at α/β KC-to-MBON-γ1pedc synapses (Figs. 10B and 10C). Pairing of α/β KC activation and BAY 41-2272 injection induced robust LTP (63.0 ± 25.0 % potentiation after 16-17 min; n = 5) with a concurrent decrease of PPR (42.3 ± 3.4 % decrease), which slowly developed over the period of ∼10 min (Figs. 10D and 10E), again replicating the observation in γ KCs. The requirement of multiple rounds of pairing was also common; one-time pairing did not induce LTP (Figs. 10F and 10 G). Taken together, γ and α/β KCs may share similar induction mechanisms of cyclic nucleotide-induced synaptic plasticity but exhibit distinct durations specifically for cAMP-dependent plasticity.

**Figure 10.**
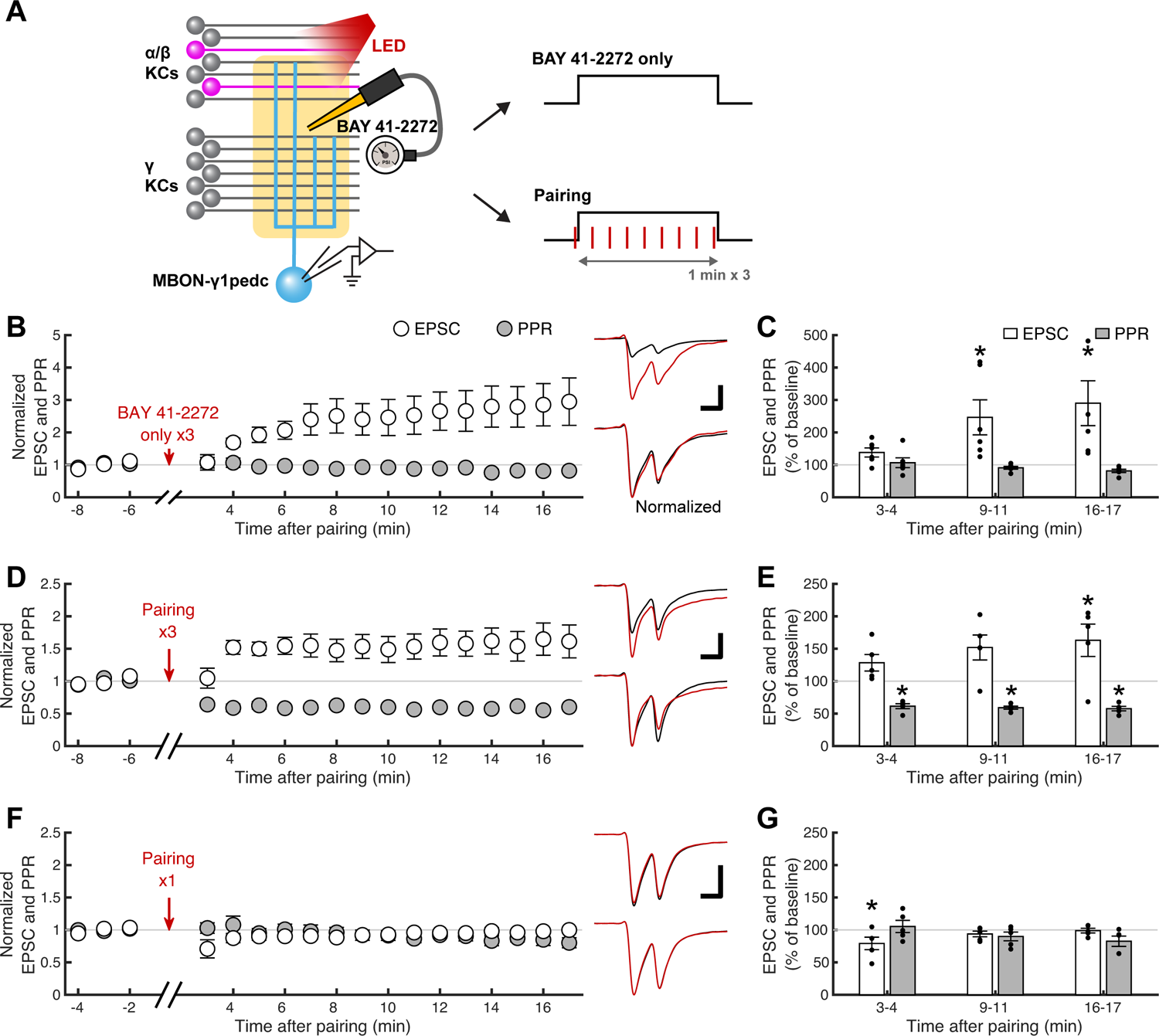
Simultaneous activation of cGMP pathway and **α**/**β**KCs induces presynaptic LTP A, a schematic of the experiments. BAY 41-2272 was focally applied to the dendritic region of the MBON-γ1pedc via an injection pipette while measuring optogenetically evoked α/β KC-to-MBON-γ1pedc EPSCs. B, first EPSC amplitudes (open circles, mean ± SEM; n = 6) and PPRs (filled circles) plotted against time after the end of the three repeats of 1-min injection of BAY 41-2272 (100 µM). The data were normalized to the average of a 3-min baseline recorded before pairing. Upper right traces show overlaid representative EPSCs sampled before (at −6 min; black) and after (at 17 min; red) injection. Horizontal and vertical scale bars in this and the other panels indicate 300 ms and 100 pA, respectively. Lower right traces show the same EPSCs normalized with the first EPSC amplitude. C, quantification of the data shown in B at early, middle and late periods after injection (mean ± SEM). Black dots indicate data from individual cells. BAY 41-2272 injection alone a delayed potentiation of first EPSC amplitudes (open bars) without coherent changes in PPRs (filled bars). P values for EPSCs are (from left to right), 0.0669, 0.000273 and < 10^−4^ (Dunnett’s multiple comparisons test following repeated measures one-way ANOVA, p < 10^−4^), and for PPRs, 0.791, 0.750 and 0.294 (repeated measures one-way ANOVA, p = 0.147). D, same as B, but BAY 41-2272 injection was paired with α/β KC activation (n = 5). E, quantification of the data shown in D. pairing of BAY 41-2272 and α/β KC activation potentiated first EPSC amplitudes and decreased PPRs at the later time points. P values for EPSCs are (from left to right) 0.488, 0.0605, and 0.0360 (Dunnett’s multiple comparisons test following repeated measures one-way ANOVA, p = 0.0502), and for PPRs, < 10^−5^, < 10^−5^ and < 10^−6^ (repeated measures one-way ANOVA, p < 10^−6^). F, same as D, but pairing was performed only once (n = 5) G, quantification of the data shown in F. One-time pairing of BAY 41-2272 and α/β KC activation was not sufficient to induce coherent changes in EPSCs or PPRs. P values for EPSCs are (from left to right) 0.0370, 0.784, and 1.00 (Dunnett’s multiple comparisons test following repeated measures one-way ANOVA, p = 0.0463), and for PPRs, 0.955, 0.380 and 0.0822 (repeated measures one-way ANOVA, p = 0.0549).

## Discussion

In many species, brain areas and cell types, activation of the cAMP/PKA pathway has been almost exclusively implicated in potentiation rather than depression of synapses in the context of synaptic plasticity. In this study, we provide direct evidence that the output synapse of the *Drosophila* MB is a rare, if not the only, exception where the sign of cAMP-induced plasticity is inverted. Our results show that potentiation is instead mediated by cGMP pathway. Against the prevailing working model, increase in neither of the cyclic nucleotides was sufficient to induce presynaptic plasticity; it additionally required simultaneous neuronal activity. Our experimental design also allowed for a separate interrogation of synaptic plasticity exhibited by different presynaptic cell types and uncovered similar but distinct properties.

Like in many sensory cortical areas, KCs show sparse sensory representations (Turner *et al*., 2008; Honegger *et al*., 2011). For this representation format to benefit the stimulus specificity of learning (Field, 1994; Olshausen & Field, 2004), the effect of neuromodulation must be restricted to the small fraction of synapses participating the sensory representation (Manoim *et al*., 2022). This requires synaptic plasticity to be induced only when neuromodulatory input coincides with synaptic activity. Calcium/calmodulin-activated AC has been long postulated as a molecular basis for such coincidence detection in multiple organisms because of its dual sensitivity to calcium influx triggered by neuronal activity and G protein signaling triggered by neuromodulatory input (Mons *et al*., 1999; Heisenberg, 2003). In *Drosophila* MB, multiple studies have indeed observed synergistic action of KC activity and DAN activation (or bath-applied dopamine) on the cAMP/PKA pathway in the KC axons (Tomchik & Davis, 2009; Gervasi *et al*., 2010; Handler *et al*., 2019). However, these and other studies also showed that DAN activation or dopamine application alone can induce considerable increase in cAMP level (Tomchik & Davis, 2009; Boto *et al*., 2014; Handler *et al*., 2019; Manoim *et al*., 2022). Our data also firmly support this observation because dopamine induced as large cAMP increase as forskolin. Moreover, DAN activity is strongly modulated by the animal’s instantaneous locomotion (Cohn *et al*., 2015; Siju *et al*., 2020; Zolin *et al*., 2021; Marquis & Wilson, 2022).

Thus, the resulting “aberrant” fluctuation of the cAMP level may prevent it from being a faithful biochemical reporter of coincidence. A recent study squarely challenged the role of cAMP as a coincidence reporter by showing that odor-electric shock pairing evoked a similar degree of cAMP elevation in the KC axons regardless of their responsiveness to the odor (Abe *et al*., 2023). Our study also supports this idea by showing the lack of synergistic action of dopamine and KC activity on cAMP synthesis. Furthermore, we did not observe cAMP increase in response to KC activation alone either. These results may seem contradictory to previous studies that reported clear cAMP increase triggered by KC activation (Tomchik & Davis, 2009; Handler *et al*., 2019; Abe *et al*., 2023) and supra-additive effect with dopamine (Tomchik & Davis, 2009; Gervasi *et al*., 2010; Handler *et al*., 2019). However, these studies used either odor presentation or acetylcholine application to stimulate KCs, which lead to cAMP increase in KCs via secondary or parallel DAN activation (Abe *et al*., 2023). In contrast, in our experiments, we optogenetically activated ∼3-7% of KCs, which is roughly equivalent to the fraction reliably responsive to a given odor (Honegger *et al*., 2011). The absence of cAMP response to KC activation and synergistic action with dopamine suggest that cAMP cannot be a reliable coincidence reporter.

The fact that cAMP does not specifically report the coincidence of dopamine input and KC activation suggests that it would be problematic if cAMP increase were sufficient to induce synaptic plasticity as assumed in the currently prevailing view (Heisenberg, 2003). That is, without another layer of coincidence detection, synapse specificity of plasticity would be compromised. Our results indicate the existence of such a mechanism. Activation of AC alone by focal application of forskolin at the synaptic site failed to induce LTD. Forskolin injection at a high concentration (100 µM) did induce LTD, but this LTD did not accompany a change in PPR. We speculate that excessive concentration of forskolin may have recruited AC in the MBON to induce postsynaptic LTD. This idea is supported by the fact that the MBON has a much lower level of AC expression compared to KCs (Fig. 4). However, the lack of change in PPR alone is not enough to specify the site of the plasticity. In general, PPR may not change after the number of release sites is altered. Conversely, PPR could change due to postsynaptic factors such as receptor desensitization (Zucker & Regehr, 2002). We were unable to analyze the miniature EPSCs, the size of which could have provided more mechanistic insight, due their small size.

Regardless of the origin of the plasticity, we can firmly conclude that the LTD induced by high concentration of forskolin is qualitatively distinct from that induced by KC-dopamine pairing because only the latter showed clear increase in PPR. In contrast to our results, a recent study reported that forskolin treatment (100 µM) alone is sufficient to induce suppression of acetylcholine release from KCs (Abe *et al*., 2023). However, this study used bath application of forskolin, and acetylcholine release was evoked by an odor. Thus, the observed effect could be the result of forskolin’s action on any part of the upstream circuit of the KCs. Moreover, they observed that the depression induced by forskolin quickly disappeared after washing out forskolin, suggesting that it is not LTD. In contrast, we focally applied forskolin only to a limited area of the MB lobes. In this small area, the resident MBON and DAN show much lower expression levels of AC compared to KCs (Fig. 4), making it unlikely that the observed effect of forskolin is mediated by non-KC cell types. In fact, the LTD induced by KC-forskolin pairing was resistant to SCH 23390, ruling out the involvement of secondary effect via potential facilitation of dopamine release. Moreover, the depression induced by KC-dopamine pairing lasted for at least 30 min after forskolin was washed out from the area, and the cAMP level returned to the baseline. We were able to replicate the LTD induced by KC-dopamine pairing only when focal forskolin application was paired with KC activation. Taken together, we propose a model that the convergence point of the signal triggered by KC activity and that by dopamine input resides somewhere downstream of the cAMP synthesis. This view and the traditional view of AC as a coincidence detector are not mutually exclusive as KC activity may have a dual role at both convergence points, although our cAMP imaging did not find the evidence for the role of AC as a coincidence detector. This potentially double-layered mechanism of coincidence detection could help secure the synapse specificity of plasticity (hence stimulus specificity of learning) and prevent the synapses from undergoing plasticity every time the cAMP level is affected by ongoing DAN activity.

Our study directly demonstrates that presynaptic cAMP increase leads to LTD, not LTP, which is in stark contrast to many other systems. In the example of *Aplysia*, serotonin-induced activation of the cAMP/PKA pathway in a siphon sensory neuron leads to phosphorylation and inactivation of presynaptic potassium channels, which facilitates the transmitter release on the motor neuron (Kandel, 2001). In hippocampal CA3, forskolin can mimic the presynaptic LTP induced by tetanic stimulation of mossy fibers, which is blocked by Rp-cAMPS (Huang *et al*., 1994).

Conversely, low-frequency stimulation of mossy fibers induces presynaptic LTD by inactivating AC via metabotropic glutamate receptors (Tzounopoulos *et al*., 1998). PKA is considered by far the most major molecule that mediates LTP/LTD at the mossy fiber terminals, and the target molecules include many active zone and vesicular proteins such as Rab3a, synapsin, RIM1a, synaptotagmin, and tomosyn (Shahoha *et al*., 2022). Some of these proteins are also implicated in memory defect in *Drosophila* (Knapek *et al*., 2010; Chen *et al*., 2011; Niewalda *et al*., 2015). Our results indicate that PKA is essential for LTD induction. Thus, to understand why the direction of cAMP-induced plasticity is opposite in this particular system, it would be important to identify the molecular targets of PKA and the detailed mechanism of the downstream convergence with neuronal activity.

Bidirectional synaptic plasticity has been reported in KC-to-MBON-γ4 synapses, where the direction of the plasticity is determined by the temporal order of KC and DAN activation (Handler *et al*., 2019). DAN activation in the absence of KC activity is also reported to strengthen the MBON response (Cohn *et al*., 2015; Berry *et al*., 2018). Recently, it was shown that presynaptic boutons that show weak calcium responses to an odor tend to be potentiated after pairing the odor with DAN activation, suggesting a complex non-linear relationship between presynaptic calcium signal and the outcome of dopamine-induced plasticity (Davidson *et al*., 2023). Our results demonstrate the presence of yet another format of synaptic potentiation mediated by cGMP. The direction of the plasticity, higher threshold for plasticity induction, and slow kinetics of plasticity development we observed match the expectation from the behavioral study that identified NO as a cotramsmitter of a subset of DANs (Aso *et al*., 2019). Since the DANs expressing NO synthase are paired up with the MBONs implicated in short-term memory, it has been proposed that NO-induced plasticity antagonizes dopamine-induced plasticity to shorten the memory retention time. However, it is not clear whether these two plasticity pathways target the same presynaptic machinery to change the synaptic strength or exist in parallel. Just like the cAMP pathway, the cGMP pathway also needs simultaneous neuronal activity to trigger the presynaptic plasticity. To understand the detailed interaction of these two pathways, the downstream molecular target needs to be identified. Recent study reported that NO-dependent cGMP signaling can trigger transcriptional changes in KCs, which are essential for forgetting of memory at 6 hours after training (Takakura *et al*., 2023). Thus, the antagonistic relationship between cAMP and cGMP second-messenger pathways controls memory acquisition and retention over multiple timescales spanning minutes to hours to days.

By taking advantage of our experimental design that allows for subtype-specific activation of KCs, we showed that the synapses made by α/β KCs on MBON-γ1pedc display shorter dopamine-induced LTD compared to the ones made by γ KCs on the same postsynaptic neuron. This result may be somewhat unexpected because the output of α/β KCs is generally considered to be important for long-term memory retrieval (Isabel *et al*., 2004; Krashes & Waddell, 2008; Trannoy *et al*., 2011; Huang *et al*., 2013). It is unlikely, however, that the observed difference between α/β KCs and γ KCs is caused by incomplete diffusion of the injected reagents because we confirmed that the signal of Texas Red-conjugated dextran infused in the injection pipette covers the entire γ1pedc region. In addition, the duration of LTD induced by pairing of α/β KC activation with 10 µM forskolin did not change even if we used 100 µM forskolin. Furthermore, the time course of cGMP-induced LTP was similar between α/β KCs and γ KCs. Thus, we conclude that the difference reflects the different properties of the synapses. It is possible that properties of synapses in the pedunculus region, which is an uncommon area for KCs to make synapses with MBONs, are somewhat different from those of typical synapses in the MB lobes. Alternatively, it is also possible that the plasticity properties observed at α/β KC-to-MBON-γ1pedc synapses generally apply to all synapses made by α/β KCs. Transcription levels of some of the PKA isoforms are markedly lower in α/β KCs than in γ KCs (Fig. 4), which might explain the difference. Importantly, the study using *in vivo* pairing of odor presentation and optogenetic DAN activation revealed that induction of LTD in MBON-α2sc took longer pairing than that in MBON-γ1 (Hige *et al*., 2015). Thus, it is possible that the shorter LTD in α/β KC-to-MBON-γ1pedc synapses reflects higher threshold of plasticity induction in α/β KCs. In this study, we did not explore a wide range of parameters of pairing protocols; we simply followed the typical pairing durations used in typical behavior studies. It requires recording from other types of MBONs with various pairing parameters to discriminate these potential scenarios.

For a long time, the learning and memory field in *Drosophila* has been strongly driven by behavioral genetics studies, which successfully made links between key molecules and behavioral defects. However, the link between those molecules and their potential roles in synaptic plasticity has not been extensively tested. In the rodent learning and memory field, a series of experiments using slice physiology provided countless key insights into the molecular and physiological basis of long-term synaptic plasticity. Our *ex vivo* system developed in this study provides an equivalent platform to interrogate the molecular machinery of synaptic plasticity in this important model organism.

## Data availability statement

All original data generated in this work are fully available to be shared upon reasonable request to the corresponding author.

## Author contributions

DY: Data curation, Formal analysis, Investigation, Writing – reviewing and editing; AMD: Data curation, Formal analysis, Investigation, Visualization, Writing – original draft, Writing – reviewing and editing; TH: Conceptualization, Formal analysis, Funding acquisition, Methodology, Project administration, Supervision, Validation, Visualization, Writing – original draft, Writing – reviewing and editing.

## Acknowledgements

We thank Yoshinori Aso and members of the Hige lab for valuable comments on the manuscript. We also thank Wanhe Li and Yulong Li for generously sharing the G-Flamp1 flies. Stocks obtained from FlyLight Split-GAL4 Driver Collection (Janelia, HHMI) and the Bloomington Drosophila Stock Center (NIH P40OD018537) were used in this study.

